# Mycobacterial formation of intracellular lipid inclusions is a dynamic process associated with rapid replication

**DOI:** 10.1101/2023.08.10.552809

**Authors:** DM Fines, D Schichnes, M Knight, A Anaya-Sanchez, NTT Thuong, J Cox, SA Stanley

## Abstract

Intracellular lipid inclusions (ILI) are triacylglyceride rich organelles produced by mycobacteria thought to serve as energy reservoirs. It is believed that ILI are formed as a result of a *dosR* mediated transition from replicative growth to non-replicating persistence (NRP). ILI rich *Mycobacterium tuberculosis* (Mtb) bacilli have been reported during infection and in sputum, establishing their importance in Mtb pathogenesis. Studies conducted in mycobacteria such as *Mycobacterium smegmatis, Mycobacterium abscessus,* or lab Mtb strains have demonstrated ILI formation in the presence of hypoxic, nitric oxide, nutrient limitation, or low nitrogen stress, conditions believed to emulate the host environment within which Mtb resides. Here, we show that *M. marinum* and clinical Mtb isolates make ILI during active replication in axenic culture independent of environmental stressors. By tracking ILI formation dynamics we demonstrate that ILI are quickly formed in the presence of fresh media or exogenous fatty acids but are rapidly depleted while bacteria are still actively replicating. We also show that the cell envelope is an alternate site for neutral lipid accumulation observed during stationary phase. In addition, we screen a panel of 60 clinical isolates and observe variation in ILI production during early log phase growth between and among Mtb lineages. Finally, we show that *dosR* expression level does not strictly correlate with ILI accumulation in fresh clinical isolates. Taken together, our data provide evidence of an active ILI formation pathway in replicating mycobacteria cultured in the absence of stressors, suggesting a decoupling of ILI formation from NRP.

## Introduction

*Mycobacteria* are a genus of bacteria that contain numerous human pathogens, the most prominent being *Mycobacterium tuberculosis* (Mtb), a deadly pathogen that causes over 1.5 million deaths annually [1]. It is estimated that up to 25% of the global population has been infected by Mtb, with many infected individuals harboring latent disease [2]. A major replicative niche of Mtb are host macrophages, cells that normally function to eliminate invading microbes. Understanding how Mtb survives and grows within macrophages is key to understanding its success as a pathogen. Lipid metabolism is central to Mtb’s ability to growth within macrophages [3]. Mce1, a transporter found in the cytoplasmic membrane of mycobacteria, is essential for importing long chain fatty acids from the environment, including fatty acids obtained from host macrophages during infection [4]. It is assumed that Mtb utilizes host lipids as a carbon source to fuel intracellular growth. Mtb is also thought to exploit lipids at late stages of infection in the lipid rich environment of host necrotic lesions [6]. Mce1 is essential for growth in macrophages and during murine infection, demonstrating that the capacity to import lipids is a critical component of Mtb pathogenesis [4], [5].

During macrophage infection Mtb accumulates cytosolic lipid bodies, organelles consisting of a triacyglyceride (TAG) rich core bounded by a phospholipid monolayer [7]–[10]. Despite the presence of lipid bodies across the kingdoms of life they have been studied predominantly in eukaryotes, largely due to their perceived scarcity in prokaryotic species. Recent advances in eukaryotic lipid droplet biology have established that these structures are not simply inert storage depots, but actively participate in a wide variety of processes including cellular stress responses, protein storage, and energy homeostasis [11]–[15]. Studies of *Actinobacteria* and *Proteobacteria* have advanced the field’s understanding of intracellular lipid inclusions (ILI), a term reserved for prokaryotic lipid bodies [16]–[18]. Work in these organisms has shed light on environmental stimuli that lead to ILI formation and suggest that ILI play a dynamic role in bacteria. For example, ILI may play a role as a site of toxic fatty acid sequestration. Lipid analysis of TAGs synthesized by *Nocardia globerula* grown on pristane, a carbon source known to inhibit β-oxidation and growth in other bacteria, revealed the ability of this bacteria to sequester toxic oxidization-derived intermediates into the TAGs [19]. Furthermore, work in *Rhodococcus jostii* demonstrated that ILI colocalize to DNA via an intermediary protein, microorganism lipid droplet small (MLDS). In mutants lacking MLDS, ILI no longer bind to DNA and bacteria suffer greater DNA damage when exposed to UV light, suggesting ILI have a role in protecting the bacilli from genotoxic stress [20].

Work in *Mycobacteria*, members of the *Actinobacteria* phylum, has further expanded the field’s understanding of ILI. Mtb isolated from human sputum has been found to contain ILI, suggesting that ILI are a feature of Mtb during human infection [21], [22]. Interestingly, lab Mtb strains form ILI in axenic culture under conditions that induce a “dormancy” regulon mimicking non-replicating states that exist *in vivo* [23], [24]. When subjected to hypoxia, nutrient starvation, or nitric oxide stress, conditions thought to emulate the phagosome or granulomas that Mtb resides in during infection, Mtb accumulates ILI [10], [23]–[29]. Interestingly, several clinical isolates form ILI when growing in axenic culture without environmental stressors, while most laboratory strains of Mtb apparently lack ILI [30]. It is not clear whether laboratory strains have lost the ability to form ILI during axenic culture, or whether this phenotype is variable even among clinical isolates. It is also possible that specific culture conditions are required to observe ILI formation. For example, *Mycobacterium smegmatis* and *Mycobacterium abscessus*, members of nontuberculous mycobacteria (NTM), were demonstrated to form ILI when subjected to nitrogen starvation simultaneous with carbon excess [31]. It is now recognized that ILI-rich states are correlated with enhanced and sustained virulence in mycobacteria. Nitrogen deprived ILI-rich *M. abscessus* are significantly more virulent compared to ILI deplete *M. abscessus* in zebrafish embryos [31]. In addition, ILI rich states have also been linked to enhanced antibiotic resistance. Carbon metabolism studies have determined that TAG synthesis inhibits growth, and bacteria induced to form ILI exhibited heighted resistance to first line antitubercular antibiotics [32]–[34].

The current consensus in the field is that Mtb forms ILI during states of non-replicating persistence (NRP) or during times of stress [21], [29], [35]. It is widely recognized that upon gradual oxygen depletion, Mtb transitions to NRP, a transition initiated by the dormancy survival regulator DosR. Approximately 48 genes constitute the DosR regulon [36]. Among these genes is *tgs1*, the predominant triacylglyceride synthase, whose disruption leads to significant reduction in TAG levels [37]. TAG accumulation and ILI formation initiated by the DosR regulon has hence been accepted as markers of successful transition to NRP [21], [35], [38]. Yet some lines of evidence suggest that ILI formation may also be associated with replicative states of the bacteria. We recently showed that Mtb forms ILI in resting macrophages that are permissive for bacterial growth, and not in IFN-γ activated macrophages that restrict bacterial replication [39]. This finding was surprising given that IFN-γ activated macrophages produce NO, a stressor known to promote ILI formation *in vitro* [26], [40]. Indeed, although there are numerous observations related to ILI formation in mycobacteria, we know very little about the conditions under which they form, the mechanism of formation, or their function. Further, although the formation of ILI in axenic culture is thought to be a feature of L2 strains of Mtb, it is unclear whether ILI formation is linked to the fitness or virulence of this lineage. Interestingly, although ILI are thought to be the primary storage depot for TAGs in bacteria, recent reports have suggested that under specific conditions, mycobacteria export TAG to their outer membrane, though the reason remains unknown [41], [42]. Thus there remains much concerning ILI formation specifically, and TAG biology more generally, that requires elucidation.

Here, we use a laboratory strain of *Mycobacterium marinum* and clinical isolates of *M. tuberculosis* to define conditions leading to ILI formation in mycobacteria. *M. marinum* is a close relative of Mtb as shown by 16S and fatty acid profiling and shares a significant percentage of essential genes with Mtb [43]–[45]. This waterborne pathogen is commonly studied in a host-pathogen model to infect the amoeba *Dictyostelium discoideum;* in these contexts, *M. marinum* has also been shown to use host lipids to generate ILI during infection [46]. Through imaging, genetic, and biochemical assays we characterize the dynamics of *M. marinum* ILI formation in axenic broth as well as in the presence of environmental factors previously demonstrated to induce ILI formation in mycobacteria. To our surprise, we discover that *M. marinum,* unique from other NTM and lab strains of Mtb, robustly forms ILI in axenic culture. This phenomenon is independent of environmental stressors and NRP, challenging the prevailing notion in the field that ILI formation is distinctly correlated with NRP or a response to stress. We also show that *M. marinum* utilizes fatty acids from media to create ILI, and that both *M. marinum* and Mtb form ILI during early log phase and then deplete them at later stages of growth. Interestingly, using super resolution microscopy, we find that at later stages of growth both species appear to accumulate TAGs in the cell envelope, further highlighting that ILI formation is specifically associated with active replication. Finally, we show that ILI formation in stress-free axenic culture may be a feature of most clinical isolates that is lost upon laboratory cultivation.

Interestingly strains of Mtb that do not produce ILI in early log phase growth appear to instead accumulate neutral lipids in their cell envelope. Gene expression analysis in ILI high and ILI low isolates demonstrated a lack of correlation between *dosR* expression level and ILI status, contrary to what was expected. Taken together, these findings demonstrate that neither NRP nor *dosR* expression are required for ILI formation in some clinical Mtb strains, suggesting that an alternative pathway may be responsible for ILI accumulation.

## Results

### *M. marinum* produces ILIs during macrophage infection

We first tested whether *M. marinum,* like *M. tuberculosis,* forms ILI in resting macrophages. We infected murine C57BL/6 bone marrow derived macrophages (BMDM) with *M. marinum* and observed robust replication of the bacteria during infection (Fig. S1, Table S1). Despite being in an ILI depleted state immediately upon infection (Fig. 1A), WT bacteria formed ILI throughout the course of the 72- hour infection, accumulating increasing levels of ILI until the point of host cell rupture (Fig. 1B, 1C). The Mce1/LucA complex has been implicated in the import of palmitate and other long chain fatty acids, leading to the formation of ILI during host cell infection by Mtb [4], [8]. Similar to Mtb, formation of ILI formation was reduced in *mce1* mutant *M. marinum*during infection of macrophages, particularly at early timepoints (Fig. 1C, 1D). *Mce1* mutants were able to form substantial levels of ILI at later timepoints, suggesting the existence of alternate fatty acid import mechanisms or *de novo* fatty acid synthesis by *mce1* mutants (Fig. 1C). The ESX-1 type VII secretion system is required for phagosomal perforation and virulence in both *M. marinum* and *M. tuberculosis* [47], [48]. In *M. marinum,* this perforation results in liberation of the bacteria into the cytosol and actin-based motility for a fraction of the bacteria [49], [50]. The ESX-1 secretion system is also required for phagosome maturation arrest, and ESX-1 mutants have increased lysosomal residence. We next tested whether ESX-1 mutants are impacted in their ability to form ILI. We found that the *M. marinum* region of difference 1 (RD1) ESX-1 mutant was attenuated for ILI formation in macrophages compared to WT *M. marinum* (Fig. 1C, 1E). The ESX-1 mutant had a greater than 60% reduction in ILI signal at 72 hours compared to WT, an attenuation similar to the fatty acid import deficient *mce1* mutant (Fig. 1C). These data demonstrate that *M. marinum* utilizes the ESX-1 secretion system and Mce1 lipid importer to acquire fatty acids from the host to form ILI. Furthermore, the fact that *M. marinum* forms ILI during active replication in host cells is counter to the prevailing hypothesis that bacteria form ILI during conditions associated with dormancy.

**Figure 1.**
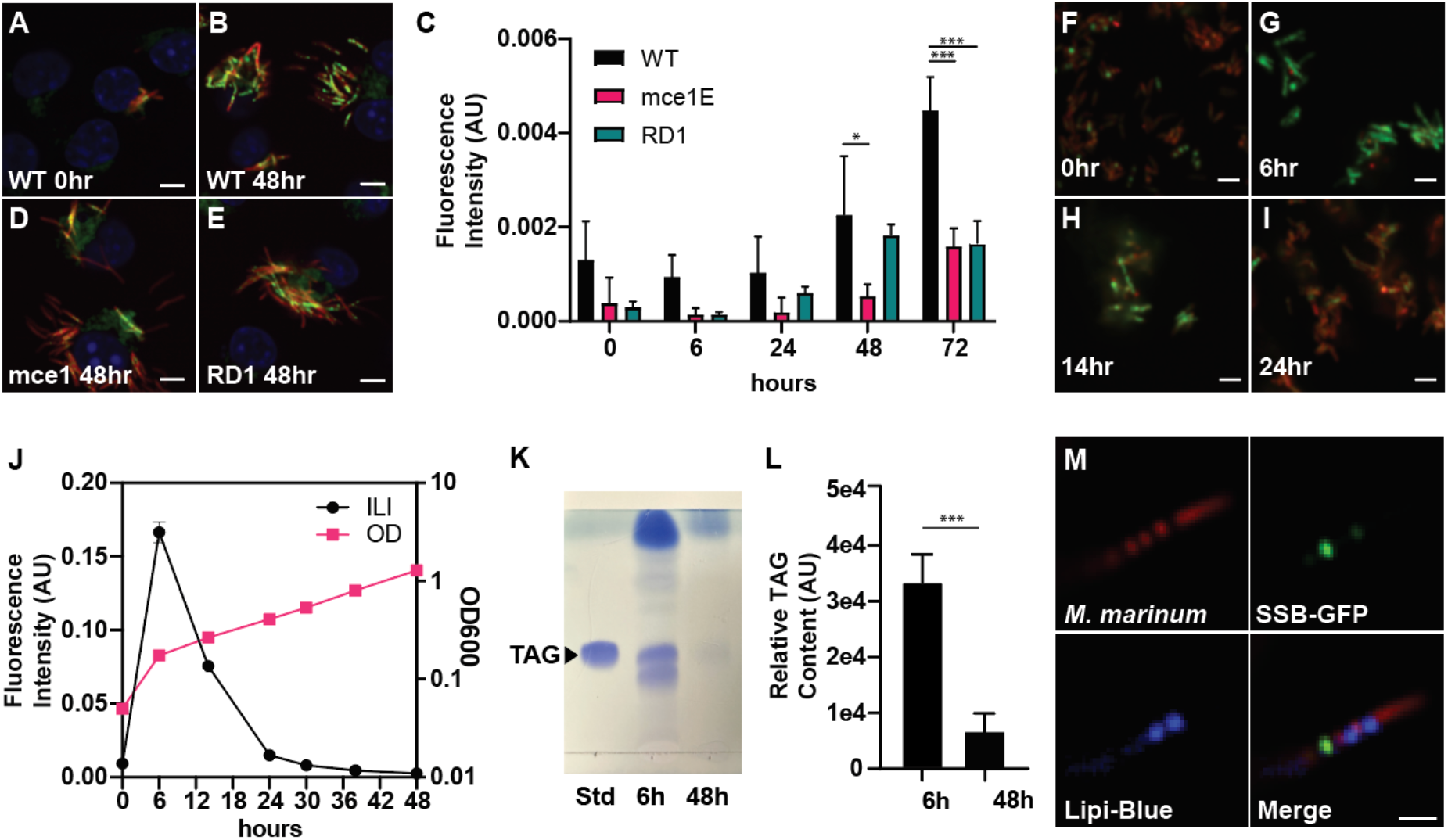
*M. marinum* forms ILIs while replicating in axenic culture. **A-E.** Confocal images of BMDM (A) immediately after infection by mCherry fluorescent WT *M. marinum* or 48 hours post infection by mCherry fluorescent (B) WT *M. marinum*, (D) *mce1* mutant *M. marinum*, or (E) *RD1* mutant *M. marinum.* Samples were stained for neutral lipids with BODIPY 493/503. Infections were at MOI=1. Scale bars represent 5 µm. (C) The integrated BODIPY 493/503 signal per bacteria was calculated with CellProfiler. Data are means ±SD (n=4). Fluorescence intensities were compared using a Student’s t-test *p<0.05, ***p<0.001. **F-J.** mCherry fluorescent WT *M. marinum* was cultured in 7H9^ADS^ and sampled at (F) 0 hours, (G) 6 hours, (H) 14 hours, and (I) 24 hours. (J) Samples collected at indicated timepoints were sampled for OD_600_ and stained for neutral lipids with BODIPY 493/503. Scale bars represent 5 µm. The integrated BODIPY 493/503 signal per bacteria was calculated with CellProfiler. Data are means ±SD (N=3). **K-L.** WT *M. marinum* was cultured in 7H9^ADS^ for 6 hours and 48 hours. Samples were visualized by (K) TLC and (L) analyzed for relative TAG abundance. Data are means ±SD (N=2). TAG band intensities were compared using a Student’s t- test ***p<0.001. **M.** SSB-GFP expressing mCherry fluorescent WT *M. marinum* was cultured for 6 hours and stained for neutral lipids with Lipi-Blue. Scale bar represents 2 µm.

### *M. marinum* forms ILIs while replicating in axenic culture

Dissecting inputs into ILI formation during infection of host cell present numerous challenges due to the complexity of the host environment, which is ill defined. We therefore tested whether *M. marinum* produces ILI during logarithmic growth in axenic culture. We grew mCherry-expressing *M. marinum* in liquid media and sampled the culture at various intervals throughout the course of bacterial growth to monitor the dynamics of ILI formation. Each sample was stained with BODIPY 493/503 to visualize neutral lipid rich puncta [4], [51]. At the time point of inoculation into a fresh culture (0h) bacterial cells lacked ILI (Fig. 1F), yet six hours post inoculation bacteria were ILI rich (Fig. 1G). ILI levels waned as the bacteria grew until a state of ILI depletion was observed at 24 hours post inoculation (Fig. 1H, 1I). Quantification of the results showed that the peak of ILI formation occurred during early log phase, and a decrease in ILI abundance correlated with a decrease in replication rate of the bacteria (Fig. 1J). By the time bacteria approached stationary phase ILI had disappeared (Fig. 1I). The fact that bacteria had low ILI levels prior to inoculation indicates that the ILI rich state at the early time points was a result of rapid neutral lipid accumulation and ILI formation while the bacteria were actively replicating in early log phase culture.

ILI in mycobacteria are classically composed primarily of TAGs [52]–[54]. To confirm that the neutral lipid puncta stained by BODIPY 493/503 were indeed ILI, we isolated lipids from cultures grown for 6 hours and 48 hours and normalized the lipid content to cell mass by OD_600_ measurements. TLC of the extracted lipids indeed confirmed that freshly cultured WT *M. marinum* has a marked increase in TAGs compared to the WT *M. marinum* cultured to a higher OD (Fig. 1K, Fig. 1L). To confirm that ILI are observed in actively replicating bacteria, we used a fluorescent reporter of single strand binding protein (SSB) that has been demonstrated to reliably identify bacteria undergoing active replication in *M. smegmatis* and *Mtb* [55]. Replicating bacteria, indicated by the SSB-GFP foci, contained Lipi-Blue stained ILI, implying replicating bacteria indeed formed ILI (Fig. 1M). As a final confirmation that *M. marinum* produces ILI during early log phase while actively replicating, we made use of phase microscopy in conjunction with a microfluidic system to capture live cell microscopy data. Phase contrast microscopy has been utilized in previous works to visualize ILI and allowed us to capture a time lapse video of *M. marinum* cells replicating [56], [57] (Fig. S2, Supplemental Video 1). Bacteria rapidly formed structures resembling ILI (Fig. S2 arrows, Supplemental Video 1). These structures occupied a significant portion of the cellular space and were capable of fusing with one another, characteristics consistent with previous findings in other prokaryotes [58].

Together, these data clearly support the finding that *M. marinum* forms ILI while actively replicating in early log phase in axenic culture. Unexpectedly, the bacteria seem to make and then deplete ILI all before reaching stationary phase, despite the fact that the media contains sufficient sources of carbon to fuel extensive bacterial growth.

### *M. marinum* formation of ILI in axenic culture requires exogenous lipids

To test if the ILI- rich status of bacteria in early log-phase growth was reflective of environmental conditions or growth phase, we grew bacteria to a timepoint at which they had depleted ILI and then replenished the culture with fresh media. Media-replenished cultures exhibited a marked increase in ILI (Fig. 2A, 2C) compared to cultures resuspended in their spent media (Fig. 2B, 2C). This result indicated ILI formation in *M. marinum* is stimulated by a component of media and is independent of bacterial growth phase. Having established that ILI formation is most robust in the presence of fresh media we sought to understand what components in the media induce *M. marinum* ILI formation. Both Mtb and *M. marinum* are known to import fatty acids during infection of host cells. We hypothesized that *M. marinum* similarly uptakes fatty acids during axenic growth and uses them to form ILI. *M. marinum* was cultured in minimal media supplemented with the sole carbon sources glycerol, acetate, stearate, and oleate (Fig. 2D-H).

**Figure 2.**
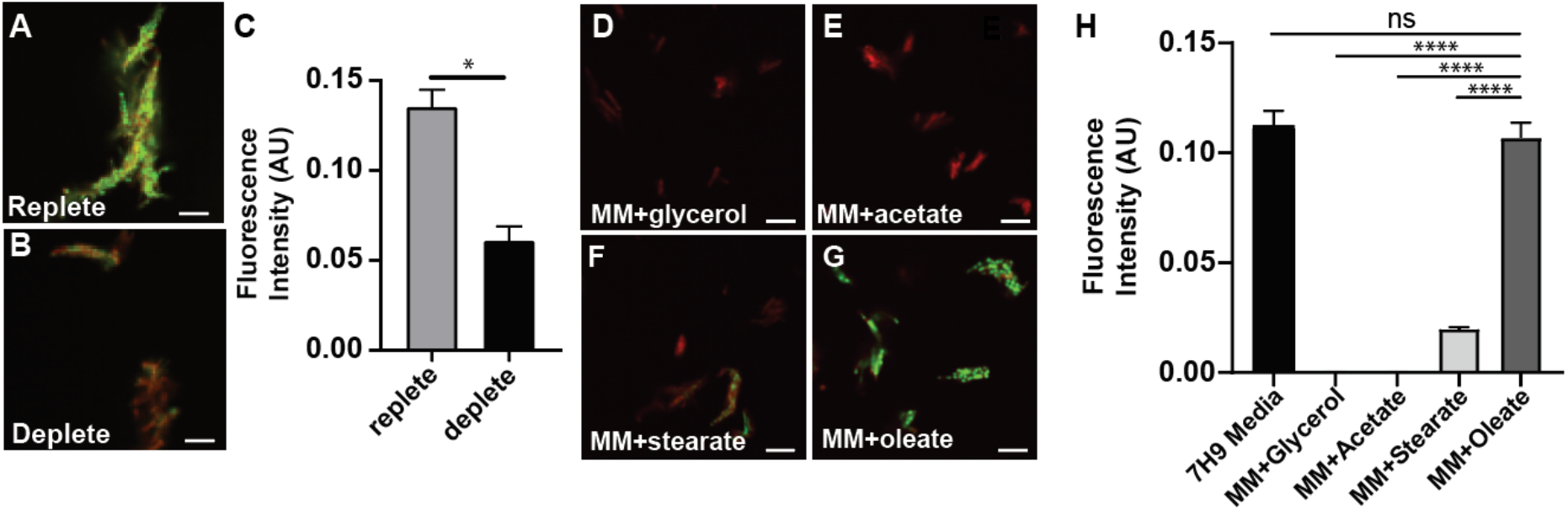
*M. marinum* formation of ILI in axenic culture requires exogenous lipids. **A-C.** mCherry fluorescent WT *M. marinum* was cultured for 6 hours and resuspended in (A) fresh, nutrient replete media or (B) the same, nutrient deplete media. Samples were collected 18 hours post resuspension and stained for neutral lipids with BODIPY 493/503. Scale bars represent 5 µm. (C) The integrated BODIPY 493/503 signal per bacteria was calculated with CellProfiler. Data are means ± SD (N=2). **D-I.** Fluorescence intensity was compared using a Student’s t-test *p<0.05. mCherry fluorescent WT *M. marinum* as cultured in (D) 7H9^ADS^ or Minimal Media supplemented with (E) glycerol, (F) acetate, (G) stearate, or (H) oleate and stained for neutral lipids with BODIPY 493/503. Scale bars represent 5 µm. (I) Integrated BODIPY 493/503 per bacteria was quantified with CellProfiler. Data are means ±SD (n=3). Fluorescence intensity was compared using a Student’s t-test ****p<0.0001.

ILI accumulation, indicated by BODIPY 493/503 staining, revealed that *M. marinum* requires the presence of long chain fatty acids in the extracellular environment to form ILI, and that abundant alternative carbon sources such as glycerol or acetate are insufficient. To test whether *mce1* is required for ILI formation, we compared WT with a *mce1* transposon mutant (Fig. S3A-C). When grown in axenic culture, the *mce1* mutant failed to produce ILI (Fig. S3B, S3C), suggesting that mce1 is a conserved component of ILI formation among *M. marinum* and Mtb. To confirm this finding, we supplemented the media with the fluorescent fatty acid BODIPY FL C_16_. The *mce1* mutant again failed to form ILI, confirming that similar to its role in Mtb, Mce1 in *M. marinum* is a critical component in the import of exogenous 16 carbon chain fatty acids (Fig. S3D-F). Thus, *M. marinum* represents a tractable system for studying ILI formation without the need for a host cell infection.

### Characterization of environmental factors contributing to *M. marinum* ILI formation

Nitrogen limitation has been well characterized to lead to ILI formation in many species, including *Rhodococcus opacus*, and the NTM *M. smegmatis* and *M. abscessus* [31], [59]. To study the consequences of a nitrogen limited environment on *M. marinum*, we cultured the bacteria in Minimal Mineral Salt Medium with standard (MSM) or low (MSM NL) amounts of nitrogen as previously described [31]. Nitrogen limitation did not lead to ILI formation in either condition, suggesting that nitrogen deprivation does not lead to a transcriptional reprogramming to promote ILI biogenesis in *M. marinum* (Fig. 3A-D). We next tested whether *M. marinum* forms ILI in in axenic culture when subjected to stressors that emulate a host environment.

**Figure 3.**
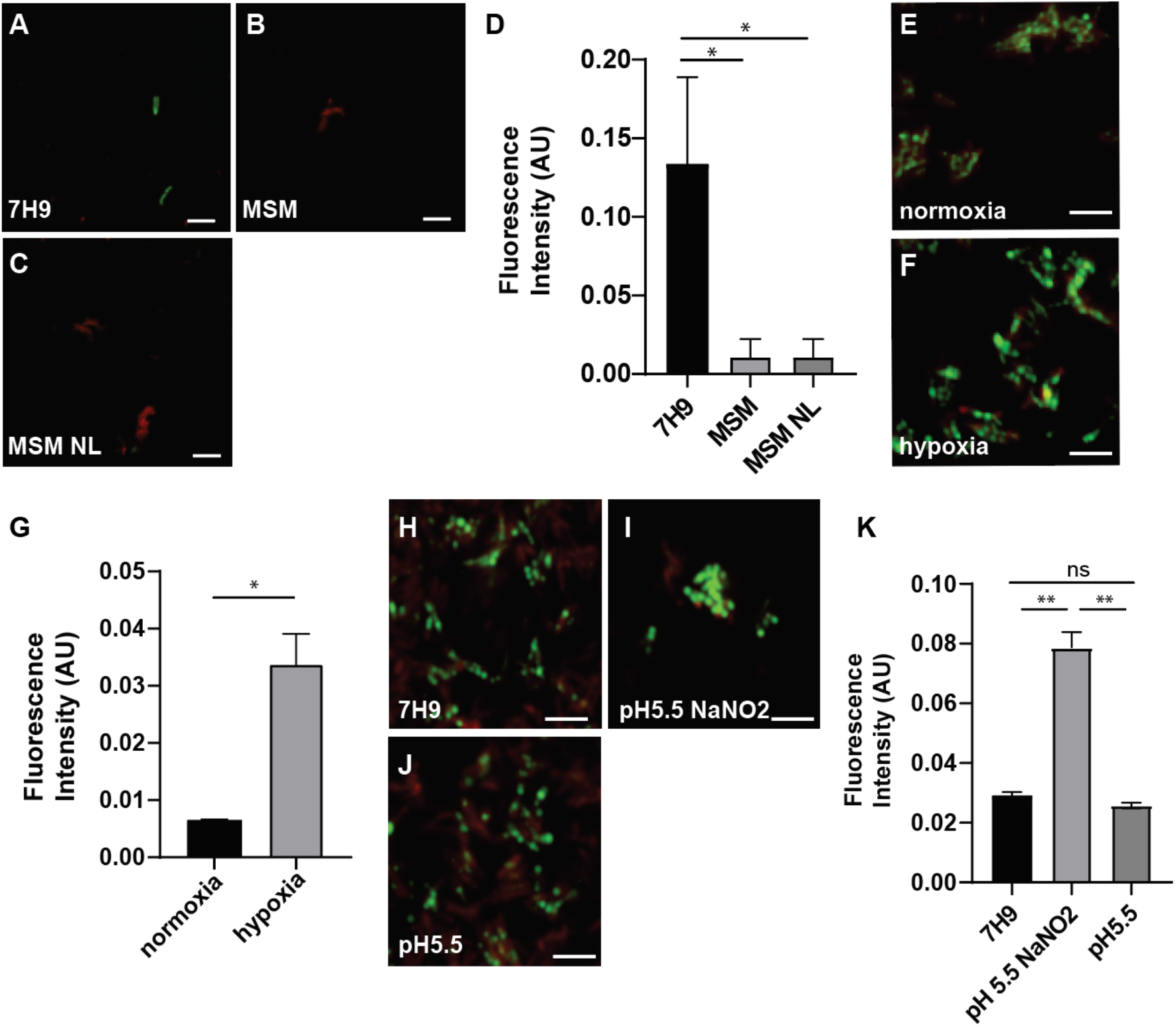
Characterization of environmental factors contributing to *M. marinum* ILI formation. **A-D.** mCherry fluorescent WT *M. marinum* strain was cultured in (A) 7H9^ADS^, (B) MSM, or (C) MSM NL media and stained with BODIPY 493/503. Scale bars represent 5 µm. (D) Integrated BODIPY 493/503 signal per bacteria was quantified with CellProfiler. Data represent means ±SD (N=3). Fluorescence intensity was compared using a Student’s t-test *p<0.05. **E-G.** mCherry fluorescent WT *M. marinum* strains were cultured in 7H9^ADS^ (E) in shaking flasks or (F) in airtight sealed inkwell bottles. Samples were collected after (E) 6 hours or (F) hypoxic conditions were reached as indicated by methylene blue and stained with BODIPY 493/503. Scale bars represent 5 µm. (G) Integrated BODIPY 493/503 signal per bacteria was quantified with CellProfiler. Data represent means ±SD (N=2). Fluorescence intensity was compared using a Student’s t-test *p<0.05. **H-K.** mCherry fluorescent WT *M. marinum* strains were cultured in (H) 7H9^ADS^, (I) 7H9^ADS^ acidified to pH 5.5 and supplemented with 1.5mM NaNO_2_, or (J) 7H9^ADS^ acidified to pH 5.5 and stained with BODIPY 493/503. Scale bars represent 5 µm. (K) Integrated BODIPY 493/503 signal per bacteria was quantified with CellProfiler. Data represent means ±SD (N=2). Fluorescence intensity was compared using a Student’s t-test **p<0.01.

Granulomas, a hallmark of infection of humans with Mtb and of zebrafish with *M. marinum*, can become hypoxic under certain conditions [24], [60], [61]. While changes in the transcriptomic landscape of *M. marinum* in hypoxic conditions has shown an increase in *tgs1* expression, it is unknown whether hypoxia stimulates ILI formation in *M. marinum* or merely TAG accumulation [62]. To examine ILI formation under hypoxic conditions, we cultured WT *M. marinum* in airtight sealed inkwell bottles supplemented with methylene blue to track oxygen depletion levels [63]. Once the cultures were confirmed to be hypoxic, we sampled the cultures and imaged ILI. We found that bacteria recovered from hypoxic cultures were highly lipid loaded, with levels four times that of a culture grown in 7H9^ADS^ for six hours (Fig. 3E-G). Other widely recognized host driven responses are the production of nitric oxide (NO) as well as the acid stress in the phagosome. Previous studies have demonstrated ILI formation, TAG accumulation, or *tgs1* expression in nitric oxide or acid stress in other mycobacteria [26]–[28], [37]. We subjected WT

*M. marinum* to acid and NO stress, culturing the bacteria in 7H9^ADS^ acidified to pH 5.5 with and without 3mM NaNO_2_ as previously described [64]. Analysis of confocal microscopy images revealed that a combination of NO and acid stress, but not acid stress alone, stimulated ILI production (Fig. 3H-K). Together, the data highlights that ILI formation is a conserved consequence of hypoxia and NO stress, while acid and nitrogen deprivation induced ILI formation is not a shared characteristic of all mycobacteria.

### *M. marinum* and Mtb can store neutral lipids in ILI or in cell membranes

To more closely examine neutral lipid staining in *M. marinum*, we turned to super resolution structured illumination microscopy (SR-SIM) [65]. Bacteria were sampled at two time points for imaging via SR-SIM. SR-SIM images revealed neutral lipid rich ILI in cultures grown for six hours, while highlighting the absence of ILI in the older cultures (Fig. 4A-F). At 48 hours we observed a shift in neutral lipid localization from ILI in the cytosolic space to the bacterial surface, suggesting that TAGs or other neutral lipids are predominantly found in the mycobacterial membrane in late log phase bacteria (Fig. 4D-F). Interestingly, BODIPY stained structures observed between some ILI at 6 hours are reminiscent of membrane bridges seen in eukaryotic cells between the endoplasmic reticulum and lipid bodies [66], [67] (Fig. 4A arrows). 3D reconstruction modeling of the SR-SIM images demonstrate that at 6 hours the BODIPY stains contoured globular structures typical of lipid bodies, while at 48 hours the BODIPY stained the perimeter of the cytosolic tdTomato signal, indicating localization to the cell membrane (Fig. 4G-H + Supplemental video 2, 3).

**Figure 4.**
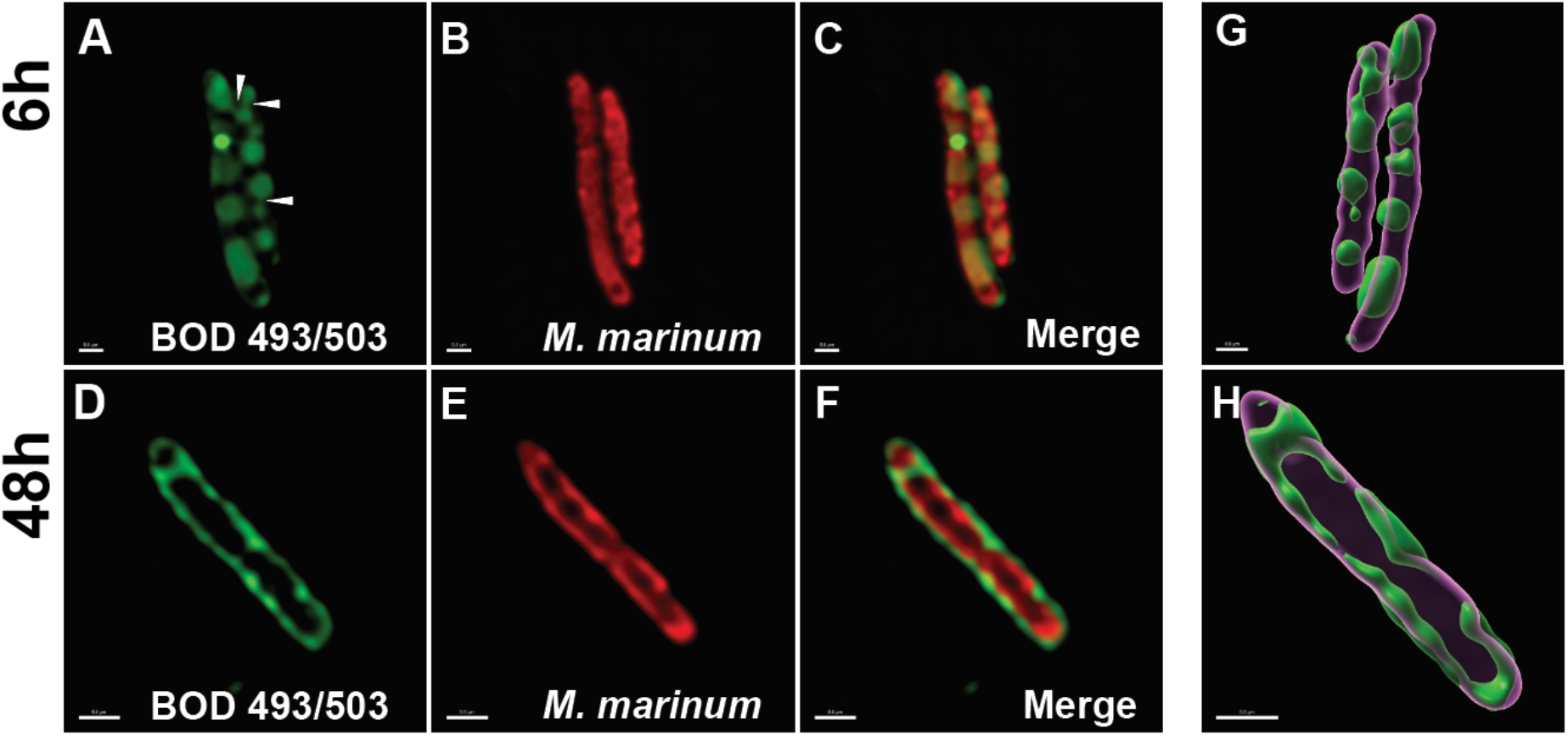
*M. marinum* can store neutral lipids in ILI or in cell membranes. **A-F.** mCherry fluorescent WT *M. marinum* strains were cultured in 7H9^ADS^ for (A-C) 6 hours or (D-F) 48 hours stained with BODIPY 493/503 and imaged via SR-SIM. White arrows in (A) indicate membrane bridge-like structures. Scale bars represent 0.5 µm. **G-H.** SR-SIM images of mCherry fluorescent WT *M. marinum* strains cultured for (G) 6 hours or (H) 48 hours were 3D modeled using Imaris. Scale bars represent 0.5 µm.

We next sought to examine ILI formation in Mtb. It has been reported that lineage 2 (L2) strains form ILI in axenic culture more readily than do other lineages [30]. Moreover, L2 strains have been associated with increased TAG accumulation [68], [69]. We compared a set of Mtb strains from lineage 4 (L4) laboratory strains (H37Rv and Erdman), as well as a L2 strain Beijing 29and a lineage 1 (L1) clinical isolate TN1401420. Cultures were back diluted to an OD_600_ of 0.05 and were sampled at 24 hours (early log) and 120 hours (late log). When stained for neutral lipids, the L2 Mtb strain showed distinct puncta associated with ILI at 24 hours, while the L1 clinical isolate and laboratory strains H37Rv and Erdman lacked these structures (Fig. 5A-D, 5F). In the L2 strain ILI were depleted by 120 hours, similar to what we observed in *M. marinum* (Fig. 5E- F). Moreover, visualization and quantification of TAG content via TLC in the strains at 24 hours revealed that the L2 strain had elevated TAG levels compared to the other strains, allowing us to confirm a link between elevated TAGs and the ILI observed by microscopy in the L2 strain (Fig. 5G, 5H). Interestingly, despite the imaging assays showing a lack of distinct ILI in the Erdman and L1 strains, we see that these strains had a significant amount of TAGs relative to H37Rv. To investigate the subcellular localization of TAGs in these strains we imaged the bacteria using SR-SIM. H37Rv showed minimal neutral lipid staining while the Erdman and L1 strains had membrane associated or discrete, amorphous staining (Fig. 5I, 5J, 5L). The L2 strain exhibited BODIPY-stained puncta as expected (Fig. 5L). 3D reconstruction modeling of the images allowed us to better visualize the lipid localization. Through this modeling we can clearly see the spherical ILI present in the L2 strain as well as the membrane associated or discrete neutral lipid localization in the other strains (Fig. 5M-P, Supplemental video 4-7). This suggests that the L2 isolate is capable of shunting synthesized TAGs into ILI while the Erdman and L1 strains, which still accumulate quantifiable TAGs, are unable to do so.

**Figure 5.**
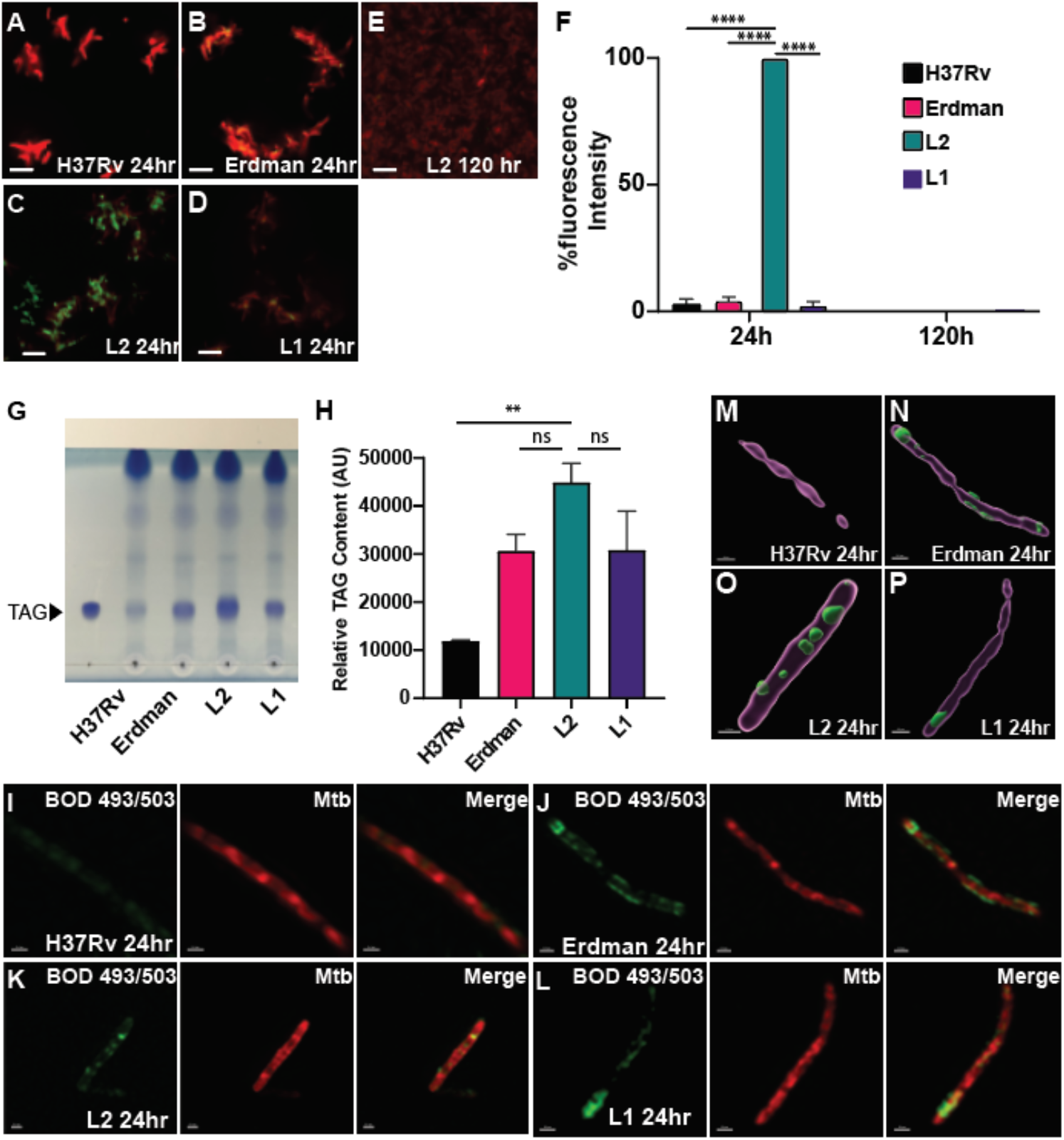
Mtb can store neutral lipids in ILI or in cell membranes. **A-F.** tdTomato fluorescent (A) H37Rv, (B) Erdman, (C) L2, and (D) L1 strains were cultured in 7H9^OADC^ for 24 hours or (E) 120 hours and stained with BODIPY 493/503. Scale bars represent 5 µm. (F) Integrated BODIPY 493/503 signal per bacteria was quantified with CellProfiler. Data represent means ±SD (N=3). Fluorescence intensity was compared using a Student’s t-test ****p<0.0001. **G-H.** H37Rv, Erdman, L2, and L1 strains were cultured for 24 hours and sampled for lipid analysis. Samples were visualized by (G) TLC and (H) analyzed for relative TAG abundance. Data are means ±SD (N=3). TAG band intensities were compared using a Student’s t-test **p<0.01. **I-L.** tdTomato fluorescent (I) H37Rv, (J) Erdman, (K) L2, and (L) L1 strains were cultured for 24 hours, stained with BODIPY 493/503, and imaged via SR-SIM. Scale bars in (I) represent 0.4 µm and (J-L) 0.5 µm. **M-P.** SR-SIM images of tdTomato fluorescent (M) H37Rv, (N) Erdman, (O) L2, and (P) L1 strains were 3D modeled using Imaris. Scale bars represent 0.5 µm.

### RNA Seq reveals transcriptional differences between ILI-producing and ILI-deficient Mtb strains

We next used RNA Seq to identify transcriptional differences that might explain the observed differences in TAG abundance and ILI formation across the Mtb strains. We compared the transcriptional signatures of the strains after 24 hours of growth post dilution. PCA analysis revealed the distinct RNA landscape of the L2 strain from the other strains as well as the clustering of the L4 lab strains H37Rv and Erdman (Fig. 6A). Differential RNA expression of genes expressed in the ILI-producing L2 strain compared to the ILI-deficient H37Rv, Erdman, and L1 strains revealed several significantly differentially expressed genes (DEGs) (Fig. 6B). Of note, the L2 strain had a significantly upregulated DosR regulon, in line with what previous groups have reported [68], [69] (Fig. 6B, Table 1, Supplementary Data 1). Indeed, GO analysis of genes upregulated in the ILI forming L2 strain compared to the other ILI deplete strains reveals significantly upregulated genes involved in response to hypoxia, oxygen levels, and stress (Table 1). The L2 strains grew similarly to the other strains, suggesting that despite the upregulated expression of stress response genes, the bacteria were not in hypoxic or stressed conditions (Fig. S4). KEGG analysis further supports these findings by clustering significantly upregulated DEGs into metabolic pathways, carbon metabolism, fatty acid metabolism, and two-component system associated pathways (Table 2). The top 20 significantly upregulated genes included several transmembrane-encoding proteins, including the membrane associated lipid transporter mycobacterial membrane protein large 4 (*mmpL4*) and its closely associated accessory protein mycobacterial membrane protein small 4 (*mmpS4*), which were not previously published to be associated with the DevR/DosR dormancy regulon [36] (Table 3). Not included in the top 20 but still significantly upregulated were the fatty acid biosynthesis associated genes *tgs1* and *tgs3* (Supplemental Data 1), which further corroborates the finding that the L2 strain had TAG rich ILI at this time point in axenic culture. Of note, a key gene involved in the Mtb NRP transcriptional landscape, isocitrate lyase *icl,* was significantly downregulated in the L2 strain. *Icl*, critical to the methylcitrate cycle and the glyoxylate shunt in Mtb, has been well characterized to be required for growth on fatty acids and survival in latent models of TB [70]– [72]. Furthermore, several lipases *lipC, lipF,* and *lipX* were also downregulated (Supplemental Data 1). The downregulation of these genes suggests that at 24 hours, the L2 strain forms ILIs but does not initiate the hydrolysis of TAGs or β-oxidation of fatty acids for growth.

**Figure 6.**
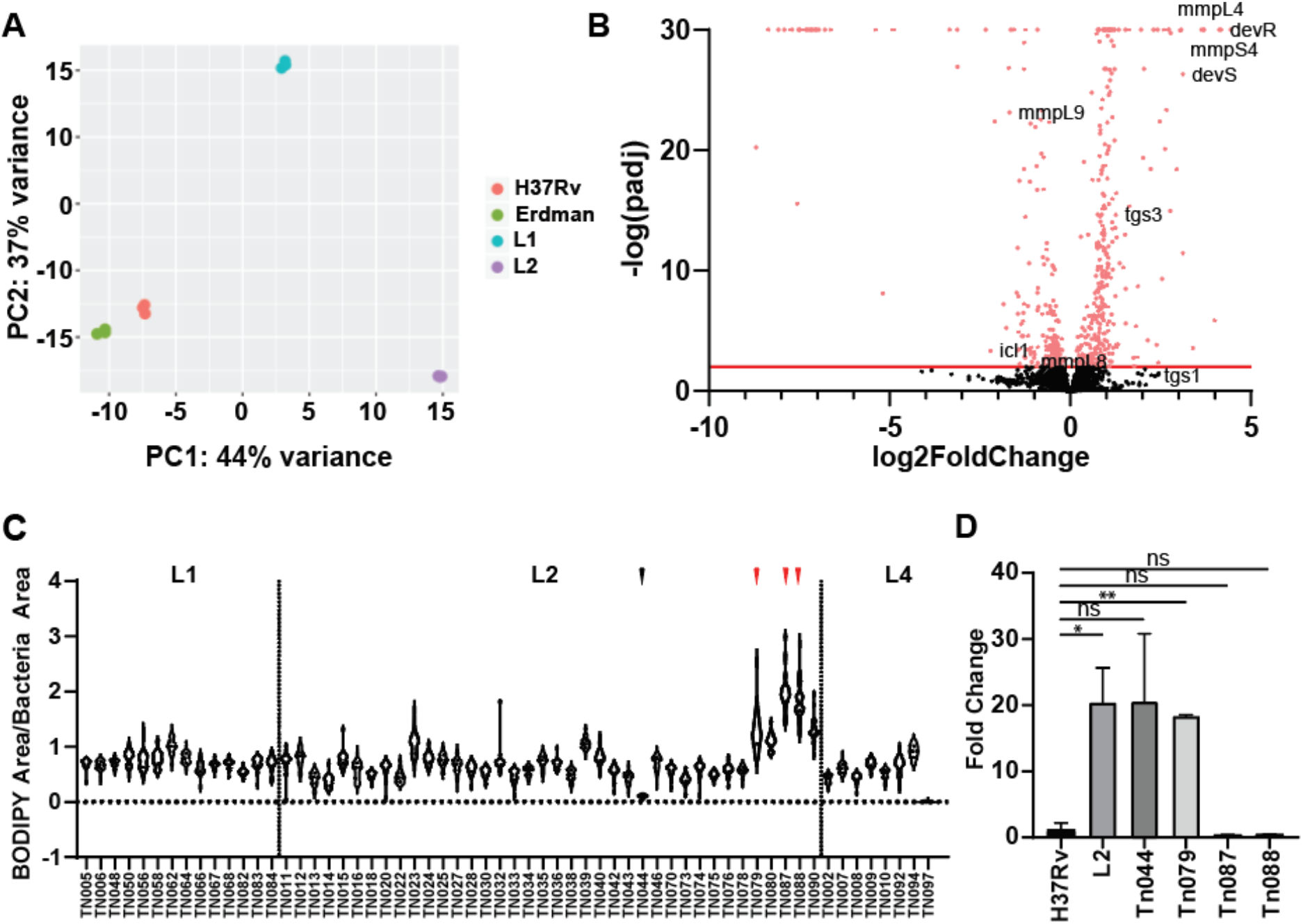
Upregulated *dosR* is not required for ILI formation in Mtb. **A.** H37Rv, Erdman, L2, and L1 strains were cultured for 24 hours and sampled RNASeq analysis. PCA analysis of the RNASeq is reflective of three biological replicates. **B.** Volcano plot of differential expression analysis comparing H37Rv, Erdman, and L1 to L2 RNA expression . The red line indicates padj (p value) = 0.05. Analysis was performed from three biological replicates. **C.** Clinical isolates were cultured for 24 hours and screened for ILI formation via BODIPY 493/503 staining. Violin plot of the integrated BODIPY 493/503 signal per bacteria is reflective of three technical replicates. Dotted lines indicate separation between the lineages. The black arrow indicates ILI low isolate TN044. Red arrows indicate ILI high isolates TN079, TN087, and TN088. **D.** Lab and clinical isolates were cultured for 24 hours and sampled for RT-qPCR analysis of *dosR* expression. Quantification was performed in reference to 16S and then normalized to H37Rv levels. Data are means ±SD (N=3). Fold change was compared using Student’s t-test *p<0.05, **p<0.01.

**Table 1.** GO analysis of significantly upregulated DEGs in the L2 strain compared to H37Rv, Erdman, and L1 strains.

| GO Name | GO ID | P-value |
| --- | --- | --- |
| GO:0009628 | response to abiotic stimulus | 1.78E-04 |
| GO:0006950 | response to stress | 3.91E-04 |
| GO:0001666 | response to hypoxia | 4.18E-04 |
| GO:0036293 | response to decreased oxygen levels | 5.70E-04 |
| GO:0070482 | response to oxygen levels | 6.94E-04 |
| GO:0000103 | sulfate assimilation | 1.63E-03 |
| GO:0010134 | sulfate assimilation via adenylyl sulfate reduction | 4.57E-03 |
| GO:0010438 | cellular response to sulfur starvation | 4.57E-03 |
| GO:0031667 | response to nutrient levels | 7.42E-03 |
| GO:0070814 | hydrogen sulfide biosynthetic process | 7.48E-03 |
| GO:0070813 | hydrogen sulfide metabolic process | 7.48E-03 |
| GO:0019419 | sulfate reduction | 7.48E-03 |
| GO:0034605 | cellular response to heat | 7.48E-03 |
| GO:0016043 | cellular component organization | 9.64E-03 |
| GO:0007154 | cell communication | 1.18E-02 |
| GO:0071840 | cellular component organization or biogenesis | 1.39E-02 |
| GO:0009408 | response to heat | 1.53E-02 |
| GO:0009267 | cellular response to starvation | 1.77E-02 |
| GO:0031669 | cellular response to nutrient levels | 1.77E-02 |
| GO:0009991 | response to extracellular stimulus | 2.00E-02 |
| GO:0000160 | phosphorelay signal transduction system | 2.13E-02 |
| GO:0042594 | response to starvation | 2.13E-02 |
| GO:0071704 | organic substance metabolic process | 2.37E-02 |
| GO:0050896 | response to stimulus | 2.41E-02 |
| GO:0009266 | response to temperature stimulus | 2.57E-02 |
| GO:0006979 | response to oxidative stress | 2.74E-02 |
| GO:0043412 | macromolecule modification | 2.94E-02 |
| GO:1901658 | glycosyl compound catabolic process | 3.07E-02 |
| GO:0022611 | dormancy process | 3.07E-02 |
| GO:0006400 | tRNA modification | 3.21E-02 |
| GO:0032502 | developmental process | 3.68E-02 |
| GO:0044238 | primary metabolic process | 4.33E-02 |
| GO:0071496 | cellular response to external stimulus | 4.53E-02 |
| GO:0031668 | cellular response to extracellular stimulus | 4.53E-02 |
| GO:0032259 | methylation | 4.85E-02 |
| GO:0016310 | phosphorylation | 4.85E-02 |

**Table 2.** KEGG analysis of significantly upregulated DEGs in the L2 strain compared to H37Rv, Erdman, and L1 strains.

| KEGG ID | KEGG Pathway |
| --- | --- |
| mtu01100 | Metabolic pathways - Mycobacterium tuberculosis H37Rv (12) |
| mtu01110 | Biosynthesis of secondary metabolites - Mycobacterium tuberculosis H37Rv (5) |
| mtu00230 | Purine metabolism - Mycobacterium tuberculosis H37Rv (4) |
| mtu01120 | Microbial metabolism in diverse environments - Mycobacterium tuberculosis H37Rv (4) |
| mtu02020 | Two-component system - Mycobacterium tuberculosis H37Rv (3) |
| mtu00920 | Sulfur metabolism - Mycobacterium tuberculosis H37Rv (3) |
| mtu01232 | Nucleotide metabolism - Mycobacterium tuberculosis H37Rv (3) |
| mtu00910 | Nitrogen metabolism - Mycobacterium tuberculosis H37Rv (2) |
| mtu00450 | Selenocompound metabolism - Mycobacterium tuberculosis H37Rv (2) |
| mtu00261 | Monobactam biosynthesis - Mycobacterium tuberculosis H37Rv (2) |
| mtu00561 | Glycerolipid metabolism - Mycobacterium tuberculosis H37Rv (1) |
| mtu03018 | RNA degradation - Mycobacterium tuberculosis H37Rv (1) |
| mtu00052 | Galactose metabolism - Mycobacterium tuberculosis H37Rv (1) |
| mtu00521 | Streptomycin biosynthesis - Mycobacterium tuberculosis H37Rv (1) |
| mtu00680 | Methane metabolism - Mycobacterium tuberculosis H37Rv (1) |
| mtu00983 | Drug metabolism - other enzymes - Mycobacterium tuberculosis H37Rv (1) |
| mtu00500 | Starch and sucrose metabolism - Mycobacterium tuberculosis H37Rv (1) |
| mtu01200 | Carbon metabolism - Mycobacterium tuberculosis H37Rv (1) |
| mtu01502 | Vancomycin resistance - Mycobacterium tuberculosis H37Rv (1) |
| mtu00470 | D-Amino acid metabolism - Mycobacterium tuberculosis H37Rv (1) |
| mtu00520 | Amino sugar and nucleotide sugar metabolism - Mycobacterium tuberculosis H37Rv (1) |
| mtu00240 | Pyrimidine metabolism - Mycobacterium tuberculosis H37Rv (1) |
| mtu00061 | Fatty acid biosynthesis - Mycobacterium tuberculosis H37Rv (1) |
| mtu01212 | Fatty acid metabolism - Mycobacterium tuberculosis H37Rv (1) |
| mtu01250 | Biosynthesis of nucleotide sugars - Mycobacterium tuberculosis H37Rv (1) |
| mtu01240 | Biosynthesis of cofactors - Mycobacterium tuberculosis H37Rv (1) |
| mtu00010 | Glycolysis / Gluconeogenesis - Mycobacterium tuberculosis H37Rv (1) |
| mtu00190 | Oxidative phosphorylation - Mycobacterium tuberculosis H37Rv (1) |

**Table 3.**
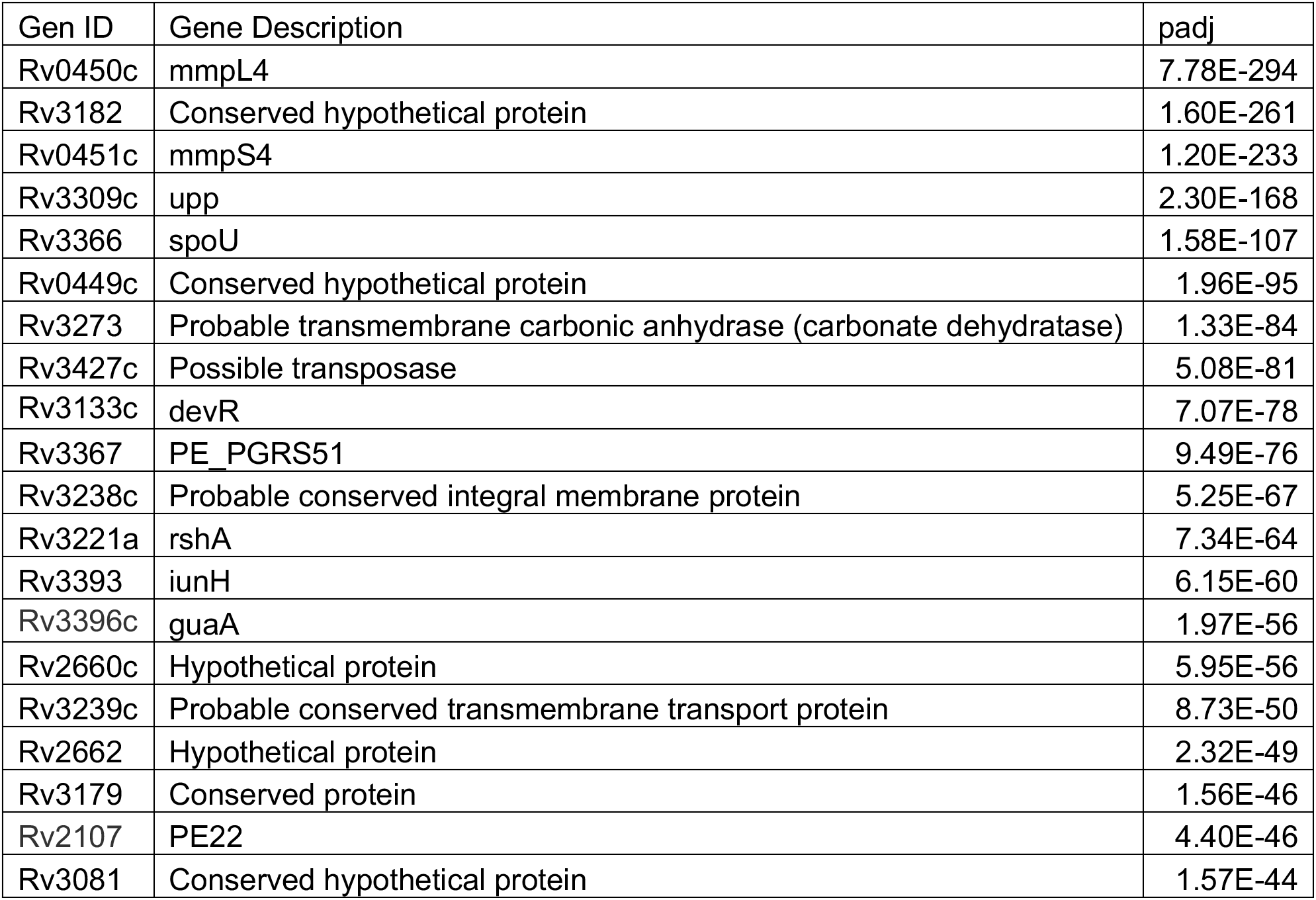
Top 20 significantly upregulated DEGs in the L2 strain compared to H37Rv, Erdman, and L1 strains.

| Gen ID | Gene Description | padj |
| --- | --- | --- |
| Rv0450c | mmpL4 | 7.78E-294 |
| Rv3182 | Conserved hypothetical protein | 1.60E-261 |
| Rv0451c | mmpS4 | 1.20E-233 |
| Rv3309c | upp | 2.30E-168 |
| Rv3366 | spoU | 1.58E-107 |
| Rv0449c | Conserved hypothetical protein | 1.96E-95 |
| Rv3273 | Probable transmembrane carbonic anhydrase (carbonate dehydratase) | 1.33E-84 |
| Rv3427c | Possible transposase | 5.08E-81 |
| Rv3133c | devR | 7.07E-78 |
| Rv3367 | PE_PGRS51 | 9.49E-76 |
| Rv3238c | Probable conserved integral membrane protein | 5.25E-67 |
| Rv3221a | rshA | 7.34E-64 |
| Rv3393 | iunH | 6.15E-60 |
| Rv3396c | guaA | 1.97E-56 |
| Rv2660c | Hypothetical protein | 5.95E-56 |
| Rv3239c | Probable conserved transmembrane transport protein | 8.73E-50 |
| Rv2662 | Hypothetical protein | 2.32E-49 |
| Rv3179 | Conserved protein | 1.56E-46 |
| Rv2107 | PE22 | 4.40E-46 |
| Rv3081 | Conserved hypothetical protein | 1.57E-44 |

### Upregulated *dosR* is not required for ILI formation in Mtb

Given the differences in ILI signal between the L1 and L2 strains, we sought to investigate if ILI variance was an inter or intra lineage trait. We thus screened 60 clinical isolates comprised of L1, L2, and L4 strains to look for any correlates of ILI signal. To our surprise, we observed both inter and intra lineage variance in neutral lipid accumulation (Fig. 6C). Even among the L2 lineages, a lineage known to accumulate TAG and producing ILI, we identified low (TN044) and high (TN079, TN087, TN088) ILI producers (Fig. 6C, black and red arrows, respectively). We can thus deduce that ILI formation may not be a trait unique to L2 strains, but rather a shared albeit variable characteristic among recent clinical isolates. The DosR regulon has been implicated to be an upstream regulator of ILI formation and has been shown to be upregulated in L2 lineages. This led us to question if there was a link between *dosR* expression and ILI formation. We turned to qPCR to examine *dosR* expression of the previously mentioned low and high ILI producers. To our surprise, we saw no correlation between *dosR* expression and ILI formation (Fig. 6D).

TN044, which exhibited quite low ILI signal, had *dosR* expression similar to the Beijing 29 and TN079 isolates. Even more surprising however was the muted expression of *dosR* in TN087 and TN088. These two isolates exhibited the highest ILI signal amongst our screened isolates yet had lower *dosR* expression than that of H37Rv. These data suggest that elevated *dosR* regulon expression is not required for ILI accumulation, separating ILI formation from a hypoxia driven dormancy regulon and hinting at another regulator or pathway distinct from dormancy regulon that drives lipid body formation.

## Discussion

ILI have long been regarded as hallmarks of NRP [21], [35], [38]. The metabolic shift that accompanies the transition from active to dormant Mtb in hypoxia is accompanied by the accumulation of lipid bodies throughout the cellular space of the bacteria. Prior to this study, the consensus in the field has been that mycobacteria such as *M. smegmatis*, *M. abscessus*, or lab Mtb strains only accumulated TAG in the presence of hypoxia, acid stress, nutrient limitation, NO, or low nitrogen stress [10], [23]–[29], [37]. While only some of these conditions have been examined by microscopy for ILI formation, it nevertheless was widely accepted that ILI and a DosR mediated TAG accumulation are hallmarks of NRP in mycobacteria [10], [21], [29], [35]. Some works however have reported findings that challenge this belief. For example, transmission electron microscopy showed that some L1 and L2 strains, but not H37Rv, formed ILI in mid log phase without the need for external stressors [30]. Furthermore, radiolabeled fatty acid-based lipid analysis showed that L2 strains accumulated TAG [68]. These reports corroborate our findings that *M. marinum* as well as Mtb clinical isolates form ILI while actively replicating independent of environmental stressors. Moreover, we made the surprising observation that ILI are formed in early log phase and are depleted as the bacteria grow. The incorporation of free fatty acids into TAG is an energetically costly process—it is difficult to imagine the benefit of TAG synthesis when the fatty acids could directly be β-oxidized to fuel cellular growth. One possiblity is that ILI serve as a means of detoxifying cytotoxic free fatty acids. Eukaryotic lipid droplets are thought to prevent lipotoxicity by integrating free fatty acids into neutral lipids such as TAGs [73]. This belief is mirrored in mycobacteria where it is speculated that fatty acid metabolism counteracts the accumulation of cholesterol and methylcitrate cycle derived cytotoxic molecules by neutralizing and rerouting these molecules into TAGs [74]. Here, we observe the accumulation of ILI and TAG in mycobacteria upon fresh media replenishment at early log or exogenous fatty acid supplementation (Fig. 2). SR-SIM and 3D reconstructive modeling of *M. marinum* highlight the accumulation of TAG in spherical ILI at the early timepoints, yet these puncta disappear at later stages and are either less prevalent or localize to the membrane (Fig. 4). In line with our observations, previous work has suggested TAG accumulates in the mycomembrane as well as at the cytoplasmic membrane[41], [42]. The initial accumulation of TAG in ILI could be a means of protection against lipotoxic stress by the bacteria wherein superfluous free fatty acids that cannot be neutralized into TAGs to be compartmentalized in the lipid bilayer are instead sequestered into ILI. At later timepoints, when excess fatty acids have been depleted from the media, there is no longer a need for upregulated TAG synthesis, and thus basal TAG levels allow for neutral lipid distribution throughout the membrane. We thus speculate that ILI accumulation is a direct consequence of fatty acid availability. This idea is further supported by our replenishment assay (Fig. 2A-C) where cultures resuspended in fresh media formed ILI while those in the spent media did not. The uncoupling of ILI formation and growth phase shown in this assay suggests that ILI formation is merely a response to exogenous free fatty acid and not necessarily due to a specific growth phase.

There remains the outstanding question of what drives the differences in ILI formation between clinical isolates. Indeed, the previously mentioned studies observed ILI in some, but not all, tested L1 and L2 strains [30]. This finding was somewhat contrary to what had been previously reported wherein TAG accumulation was found in only L2, and not L1 and L4, isolates [68]. The observed lack of TAG in L1 strains by the latter report is surprising, as our results show elevated TAG levels in an L1 strain compared to H37Rv as well as some quantifiable ILI accumulation in many L1 clinical isolates (Fig. 5G-H, Fig. 6C). The discrepancy between the findings could be explained by differences in culture conditions and captured time points. Nevertheless, the collective studies on the variance in ILI aggregation between lineages do raise the question of the genetic drivers of ILI formation. The DosR regulon regulates the transcription of many key genes in response to gradual oxygen deprivation and is thought to be the stress response pathway upstream of ILI formation [21]. Indeed, *tgs1*, the predominant triacylglycerol synthase, is one such gene whose induction is downstream of *dosR* [36]. In our study we make a surprising observation that decouples *dosR* expression from ILI formation and implicates the existence of a yet to be characterized ILI synthesis pathway. We observed that the ILI high TN087 and TN088 clinical isolates had significantly reduced *dosR* expression. In contrast, the ILI low strain TN044 had elevated *dosR* expression on par with the ILI high TN079 (Fig. 6D). Further work is being done to identify enriched SNPs associated with high ILI accumulation, as well as transcriptional profiling across a larger number of isolates to determine any genetic expression correlates to ILI level. It would be interesting to see if these investigations result in the identification of key genes directly involved in the ILI formation pathway. It is worth noting however that ILI formation in the absence of any facilitator proteins has been speculated. The composition of the lipid membrane has been theorized to be a driving force involved in the biogenesis of lipid bodies, with works on the ER membrane and in artificial lipid bodies demonstrating that dysregulated phospholipid composition and membrane surface tension alters lipid body aggregation [75]–[79]. It would be exciting to see if any of the mycobacterial differentially expressed genes or enriched SNPs were involved in membrane composition homeostasis.

Our study also highlights that although we observed ILI formation in axenic culture absent of external stressors, we found that some conditions, such as hypoxia and NO stress, led to TAG accumulation as previously reported [28], [29]. However, we failed to see ILI in *M. marinum* cultured in nitrogen limited conditions (Fig. 3A-D). It is unclear how a low nitrogen/carbon ratio environment differentially affects *M. marinum*. Unlike *M. abscessus* or *M. smegmatis*, *M. marinum* has a wide range of hosts that can serve as a replicative niche for this environmental waterborne bacterium [46], [80]. The ability to infect freshwater and marine fish, amoeba, and humans necessitates flexible adaptation to diverse environments. It is plausible that the nitrogen/carbon ratios tested in this study, despite being harsh enough to stimulate ILI formation in *M. abscessus* and *M. smegmatis*, may not have been extreme enough for *M. marinum*. The effects of an altered nitrogen/carbon ratio have yet to be tested on lab and clinical Mtb strains— these findings may provide a hint about the differences between species’ response to nitrogen deprivation. The discrepancy in response to stressors leads to speculation about the role of ILI. It has previously been thought that ILI are formed as a response to stress such as those found in the granuloma [10], [21]. Yet, as our findings show, *M. marinum* only forms ILI in some but not all previously identified stressors. Moreover, *M. marinum* and clinical Mtb isolates readily form ILI without these stressors (Fig. 1, Fig. 6C). It thus proves difficult to ascertain if ILI have a role in response to or protection from stressors. Numerous studies have attempted to link ILI formation with response to stress. For example, *tgs1* has been shown to be one of the 48 genes in the hypoxia induced DosR regulon [36]. Furthermore, several reports have demonstrated the necessity of functional *tgs* and *mce1* expression *in vivo*, suggesting ILI are critical for sustained virulence in the host [4], [5], [81]. However, while these models result in ILI loss, they also lead to a dysregulated central metabolism. Metabolomics have delineated the incorporation of fatty acids into the central carbon metabolism [82], [83]. Furthermore, studies on *icl* knockout strains have found these mutants to be attenuated *in vivo*, highlighting the importance of fatty acid catabolism [70], [72], [84]. Mtb strains with disrupted *tgs* or *mce1* expression would impact fatty acid import and TAG accumulation, resulting in a severely imbalanced central carbon flux and cause possible misinterpretations of the role of ILI. While some evidence does suggest ILI can play a role beyond the compartmentalization of energy, it is currently impossible to decouple ILI from fatty acid metabolism. It would be interesting to probe the ILI formation pathway for proteins that mediate the construction of the physical ILI structure without a major disruption of TAG synthesis or fatty acid availability.

This study also captured ILI dynamics in a previously unstudied infectious setting. Although prior work had characterized ILI in the context of *D. discoideum* infection, the culture conditions of *D. discoideum* require the bacteria to be subjected to suboptimal temperatures. We wondered if the kinetics of ILI generation and degradation seen in this host-pathogen model were representative of clinical cases, where *M. marinum* infects patient extremities at a temperature closer to its’ optimal replicative climate of 30-33°C. Interestingly, we found that BMDM infecting *M. marinum* accumulated ILI throughout the infection, unlike the accumulation and subsequent depletion of ILIs in axenic culture (Fig. 1). These ILI dynamics of ILIs in macrophage infections mirror that of *M. marinum* in *D. discodeum*, suggesting the cooler infectious climates do not seem to strongly impact ILI accumulation in *M. marinum* [46]. We saw that neither the ESX-1 mutant nor *mce1 M. marinum* mutants were able to accumulate ILI to the same degree as the WT strain. Of note, although the ESX-1 and *mce1* mutants exhibited ILI formation deficiencies, they were still able to form quantifiable amounts of ILI. Mycobacteria in the absence of LCFA are able to biosynthesize *de novo* fatty acids through *fas*, which encodes a fatty acid synthase. It is likely that despite a nutrient limited environment caused by the ESX-1 and *mce1* mutants, mutant *M. marinum* biosynthesizes free fatty acids to incorporate into TAG.

Many facets of ILI in mycobacteria remain unexplored: the specific transcriptomic and proteinaceous drivers of ILI formation, the means of nutrient acquisition by the bacteria in host cells, and the role of ILI in pathogenesis are all critical components of ILI biology that require further study. Here, we highlight that *M. marinum*, unique to other NTM, forms ILI while replicating in axenic culture. This trait is shared across L1, L2, and L4 lineages and not among lab strains of Mtb, demonstrating the importance and relevancy of using *M. marinum* as a proxy to study ILI in the clinical context. Our RNASeq results identified many potential targets for future investigation of ILI associated genes; further characterization these genes as well as examination of the differences in the membrane mosaic of mycobacteria could provide insight into the molecular machinery that dictates ILI formation. Finally, an examination of *dosR* expression and ILI accumulation highlighted a remarkable finding that ILI formation is independent of *dosR* expression. Together, our findings reveal that neither a non-replicating state nor an upregulated DosR regulon are required for ILI formation in *M. marinum* and clinical Mtb isolates. Continued characterization of this finding as well as other aspects of ILI formation are critical to understanding the Mtb dormancy state and will identify promising therapeutic targets against LTBI infection and the global TB pandemic.

## Materials and Methods

### Bacteria and Media

*M. marinum* strain M was grown in 7H9^ADS^ media (Middlebrook 7H9 Broth supplemented with 0.2% Tween-80, 0.2% Glycerol, 0.5% BSA, 0.2% dextrose, 0.085% NaCl). For single carbon source assays *M. marinum* was cultured in minimal media (0.08% NaCl, 0.04% L-asparagine, 3.5% Na_2_HPO_4_, 1.48% KH_2_PO_4_, 0.04% 5M Ammonium Iron Citrate, 0.02% 1M MgSO_4_7H_2_O, 0.05% 1mg/mL CaCl_2_, 0.01% 0.1mg/mL ZnSO_4_, 0.2% EtOH, 0.2% Tyloxapol) supplemented with either 30 mM glycerol, 30 mM sodium acetate, 200 µM stearic acid, or 200 µM oleic acid. Cultures were shaken at 100 rpm at 32°C. *M. tuberculosis* H37Rv and Erdman were grown in 7H9^OADC^ (Middlebrook 7H9 Broth supplemented with 0.04% Tween-80, 0.2% Glycerol, 10% Middlebrook OADC Enrichment). The L1 clinical isolate TB1401420 was generously provided by the Kato-Maeda lab at the University of California, San Francisco, and was cultured in 7H9^OADC^. The L2 clinical isolate was generously provided by the Darwin lab at New York University and cultured in 7H9^OADC^. All Mtb strains were cultured at 37°C and rolled at 5 rpm.

### Generation of SSB-GFP M. marinum

SSB-GFP plasmid was generously provided by the Russell lab at Cornell University. Electroporation of donor plasmids was conducted in electrocompetent *M. marinum* as previously described [85]. Transformed colonies were selected on Middlebrook 7H10 agar plates with 50 µg/mL hygromycin and cultured in 7H9^ADS^ supplemented with 50 µg/mL hygromycin.

### Fluorescent and Phase Microscopy

For imaging of bacteria in axenic culture, bacteria were cultured as described above. For fluorescent microscopy, cultures were sampled and fixed with equal amounts of 10% formalin at the noted time points. Samples were pelleted at 2850 *g* for 5 minutes, washed with equal volume 1x PBS, and resuspended with equal volume of 5uM BODIPY 493/503 in PBS for or 5 µM LipiBlue in PBS for 45-60 minutes. Samples were washed twice with 1x PBS and resuspended in a final volume of 200 µL 1x PBS before being transferred into Phenoplate 96 plates. For imaging of bacteria in host cells, samples were washed with 1x PBS and fixed in equal amounts of 10% formalin. Samples were then washed with 1x PBS and incubated with an equal volume of 5 µM BODIPY 493/503 and DAPI 1:10000 in PBS for 45-60 minutes, with a final wash in 1x PBS. All plates were imaged on an Opera Phenix High Content Screening System. Resulting images were analyzed using CellProfiler image analysis software. For live imaging, mCherry fluorescent *M. marinum* cultures were grown to OD_600_ 0.6 in 7H9^ADS^ and then back diluted to 0.05. 200 µL of back diluted cultures were loaded into CellASIC ONIX Microfluidic Plates which were then connected to a CellASIC flow chamber. Phase contrast images were taken on an AxioObserver Z1 inverted microscope at 15 minute intervals over the course of 17 hours. Super-Resolution Structured Illumination Microscopy images were obtained on a ZEISS Elyra 7 inverted microscope and 3D reconstruction modeling was performed on the Imaris Cell Imaging software.

### Lipid Extraction

*M. marinum* cultures were grown in 7H9^ADS^ media as described above. At specific time points cultures were pelleted at 2850 *g* for 5 minutes and washed 2x with Optima water. Total lipids were extracted via a 1:2 MeOH:Chloroform solution. The cultures were left rocking at room temperature for 1 hour and then spun down at 2850 *g* for 10 minutes. After the upper organic layer was transferred to a fresh tube a fresh aliquot of 1:2 MeOH:Chloroform was added to the remaining pellet. The pellet was rocked for another hour, spun at 2850 *g* for 10 minutes, and the upper organic layer was transferred to the corresponding tube. The solvent was dried under a gentle stream of nitrogen. The resulting lipid residue was resuspended in 1:2 MeOH:Chloroform.

### Thin Layer Chromatography

TLC was conducted on Hard Layer Silica Gel UNIPLATES. The solvent mixture used for the analysis of TAG was toluene:acetone 99:1 (v/v). Spots were visualized by Coomassie Blue (0.2% in 20% MeOH) for 20 minutes followed by a destaining in 20% MeOH for up to 1 hour. Imaged plates were analyzed using the FIJI imaging package.

### Replenishment Assays and BODIPY FL C_16_

*M. marinum* was cultured in 7H9^ADS^ media to mid log phase and back diluted to an OD_600_ of 0.05. The cultures were then grown for 6 hours, and then pelleted at 2850 *g* for 5 minutes. Replenished cultures were resuspended in fresh 7H9^ADS^ media while unreplenished cultures were resuspended in their original media. Samples were taken 18 hours post resuspension. For BODIPY FL C_16_ supplementation, *M. marinum* with a transposon insertion in *mce1* was cultured in 7H9^ADS^ media to mid log phase and back diluted to an OD_600_ of 0.05. The culture was grown for 48 hours. BODIPY FL C_16_ was added to a final concentration of 2.5 µM for 4 hours. The culture was washed 1x with PBS and fixed with 10% formalin. Samples were then washed 2x with PBS and imaged immediately.

### Environmental Stress Conditions

*M. marinum* cultures were grown to an OD_600_ of 0.6. For hypoxia assays, cultures were back diluted to an OD_600_ of 0.2. 20 mL of back diluted culture were placed in airtight Nalgene inkwell bottles. Methelyne blue was supplemented to track oxygen loss as previously described [63]. For acid stress, mid log cultures were washed in acidified media (classical media acidified with HCl to pH 5.5), and back diluted to OD_600_ 0.05 in acidified media. For NO stress, mid log cultures were washed in acidified media, and back diluted to OD 0.05 in acidified media supplemented with NaNO_2_ (final concentration 1.5 mM) as previously described [64]. For nitrogen limited stress, mid log cultures were washed in minimal mineral salt medium (2 g/L Na_2_HPO_4_, 1 g/L KH_2_PO_4_, 0.5 g/L NaCl, 0.2 g/L MgSO_4_, 20 mg/L CaCl_2_, and 1 g/L NH_4_Cl) or mineral salt medium nitrogen limiting (2 g/L Na_2_HPO_4_, 1 g/L KH_2_PO_4_, 0.5 g/L NaCl, 0.2 g/L MgSO_4_, 20 mg/L CaCl_2_, and 0.05 g/L NH_4_Cl), and back diluted to OD_600_ 0.05 in minimal mineral salt medium or mineral salt medium nitrogen limiting as previously described [31].

### Bacterial Infections

Bone marrow derived macrophages (BMDM) from C57BL/6 mice were seeded at 5e4 cells/ well into PhenoPlate 96 well plates 48 hours prior to infection in cell culture media (DMEM supplemented with 10% FBS, 10% CSF, and 1% glutamax). BMDM were placed in 33 °C tissue culture incubators 1 hour prior to infection to acclimate the cells to the appropriate temperature. Prior to infection, *M. marinum* was cultured in classical media to an OD_600_ of ∼0.6. Bacteria cultures were pelleted at 2850 *g* for 5 mins and washed twice with 1x PBS. A slow spin speed at 58 *g* was performed to pellet clumped bacteria. The supernatant was collected and diluted into phagocytosis media (DMEM supplemented with 5% horse serum and 5% FBS) to achieve a multiplicity of infection of 1. Bacterial suspensions were added onto cells and spun at 335 g for 10 minutes. Infected cells were washed with 1x PBS and incubated in cell culture media at 33 °C. When necessary, media changes were performed every 2 days.

### Isolation of RNA and RNA Seq

*M. tuberculosis* strains were cultured to mid log phase and back diluted in 30 mL of 7H9^OADC^ to an OD_600_ of 0.05. After 24 hours the cultures were pelleted at 2850 *g* for 5 minutes and resuspended in 1 mL Trizol. Samples were transferred to an o-ring tube with RNase and DNase free glass beads and bead beat for 30 seconds 3 times. Samples were pelleted and the supernatant was transferred to a fresh tube with 200 µL chloroform. Samples were centrifuged at 12000 *g* for 10 minutes at 4°C. Resulting aqueous phase was transferred to a fresh tube. Equal volume of 70% RNase free EtOH was added. Total RNA extraction was performed via a Qiagen RNease Mini Kit. Prepped samples were submitted to Azenta Life Sciences for RNA Sequencing and analysis.

### Screening of Clinical Isolates

Mtb strains were cultured in 10 mL of 7H9^OADC^ to mid log phase. Cultures were back diluted to an OD_600_ of 0.05 and grown for 24 hours. Samples were pelleted and fixed as described above. Fixed samples were stained with 5 µM BODIPY 493/503 and DAPI 1:10000 in PBS for 45-60 minutes, with a final wash in 1x PBS. Samples were imaged and analyzed as described above.

### RT-qPCR

*M. tuberculosis* strains were cultured to mid log phase and back diluted in 30 mL of 7H9^OADC^ to an OD_600_ of 0.05. Samples were collected 24 hours after back dilution. RNA was isolated as described above. For qPCR, cDNA was generated from 1 μg of RNA using Superscript III (Invitrogen Life Technologies) and random hexamer primers. Genes were analyzed using KAPA SYBR FAST qPCR master mix (Roche Kapa Biosystems). Each sample was analyzed in at least duplicate on a CFX96 Real-time PCR detection system (Bio-Rad). C_Q_ values were normalized to values obtained for 16S and relative changes in gene expression were calculated using the ΔΔC_Q_ method.

---

**Figure S1.**
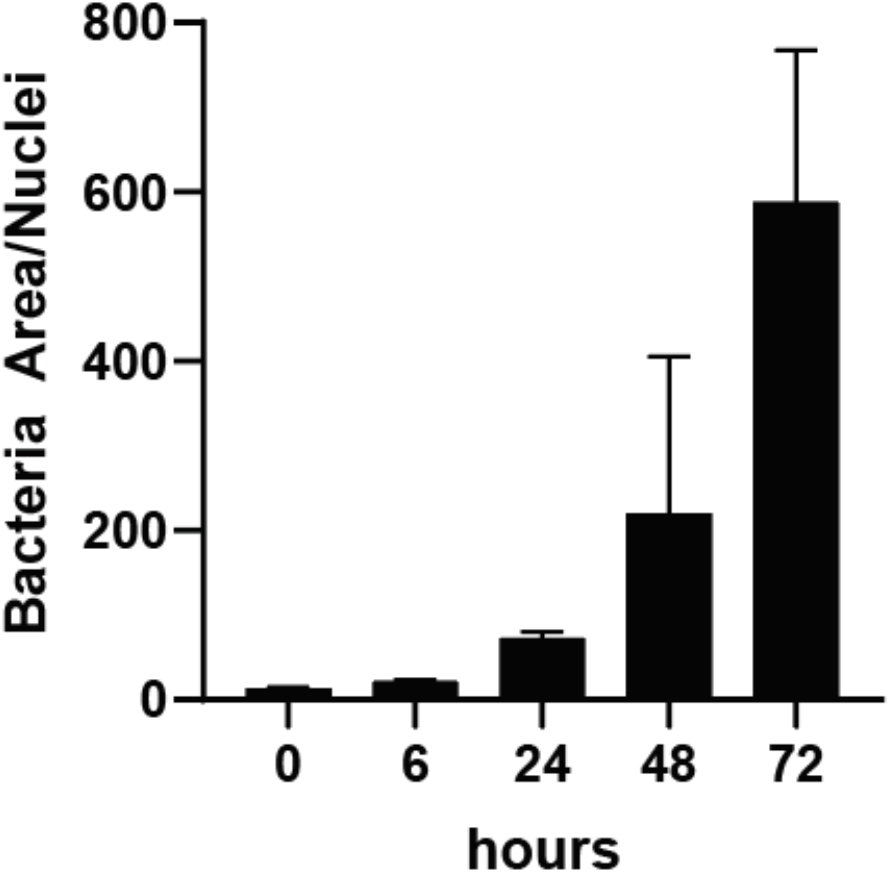
Quantified WT *M. marinum* growth in BMDM. BMDM were infected with mCherry fluorescent WT *M. marinum* at MOI=1. Samples were taken at indicated timepoints and stained with DAPI. The mCherry signal per macrophage nucleus was quantified using CellProfiler. Data represent means ±SD (n=4).

**Figure S2.**
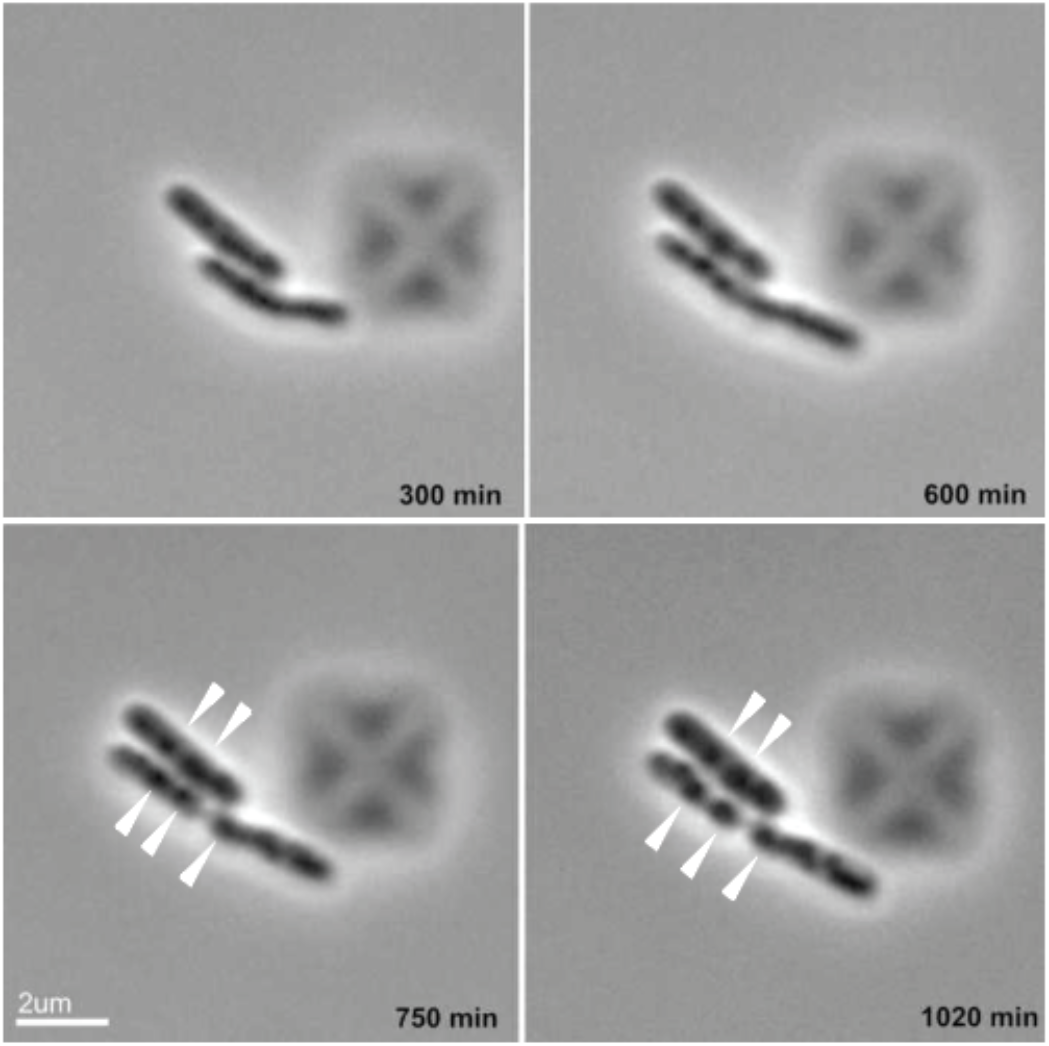
Live imaging of *M. marinum* cultured in CellASIC flow chamber. mCherry fluorescent WT *M. marinum* was cultured in a CellASIC flow chamber with 7H9^ADS^ media flowing through the system. Phase contrast images were taken every 15 minutes over the course of 17 hours. Images capture a replicating *M. marinum* bacterium with ILI (white arrows). Scale bar represents 2 µm.

**Figure S3.**
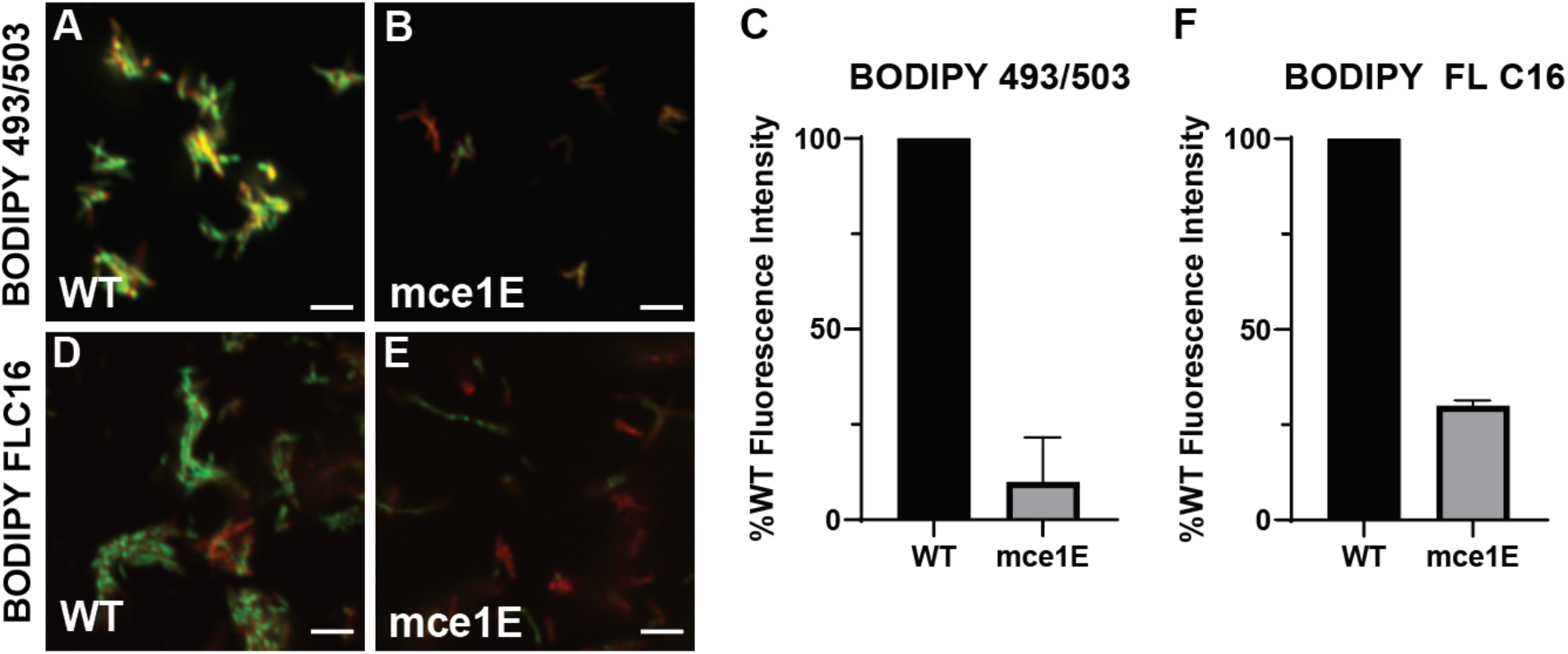
*Mce1 M. marinum* mutants are deficient for ILI formation and fatty acid import. **A-C.** mCherry fluorescent (A) WT *M. marinum* and (B) *mce1* mutant *M. marinum* were cultured for 6 hours in 7H9^ADS^ and stained with BODIPY 493/503. (C) Integrated BODIPY 493/503 signal per bacteria was quantified with CellProfiler and normalized to WT *M. marinum* signal. Data represent means ±SD (N=3). **D-F.** mCherry fluorescent (D) WT *M. marinum* and (E) *mce1* mutant *M. marinum* were cultured for 48 hours in 7H9^ADS^ and pulsed with 2.5 µM BODIPY FL C_16_ for 4 hours. Data represent means ±SD (N=2).

**Figure S4.**
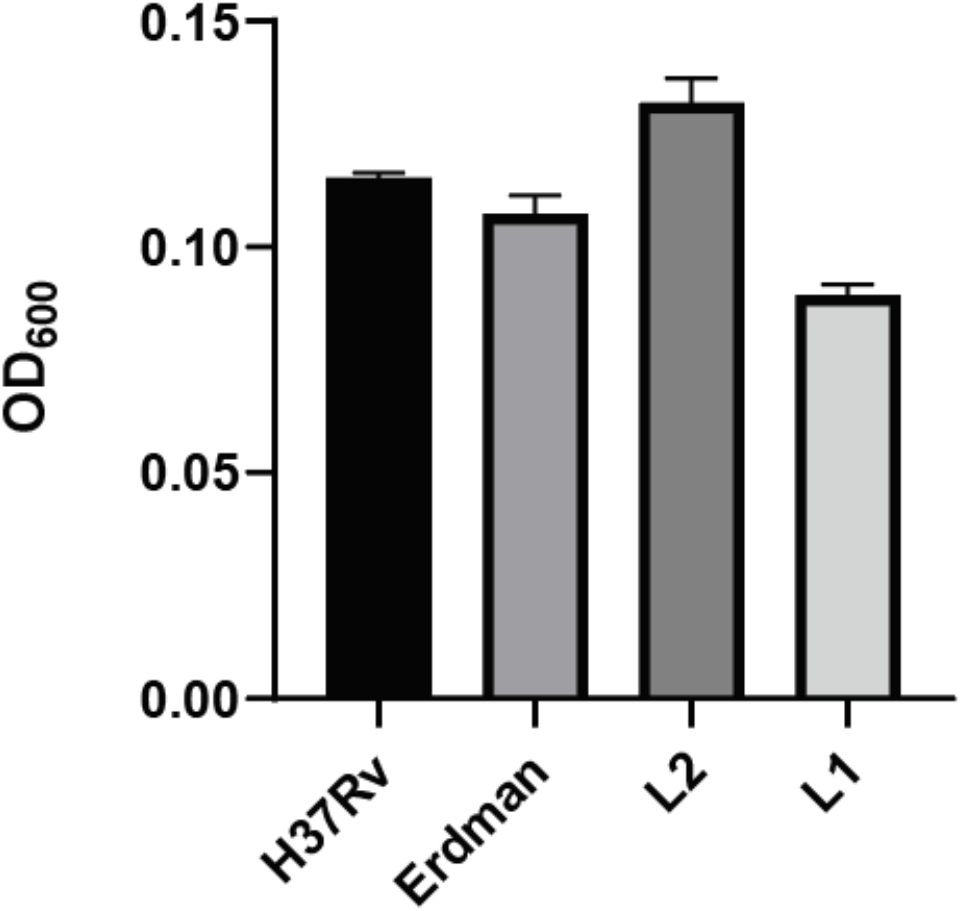
OD_600_ of Mtb cultures at time of sampling. H37Rv, Erdman, L2, and L1 strains were cultured in 7H9^OADC^ and back diluted to OD_600_ 0.05. Cultures were sampled for OD_600_ readings 24 hours post inoculation. Data represent means ±SD (N=3).

**Supplemental Table 1.**
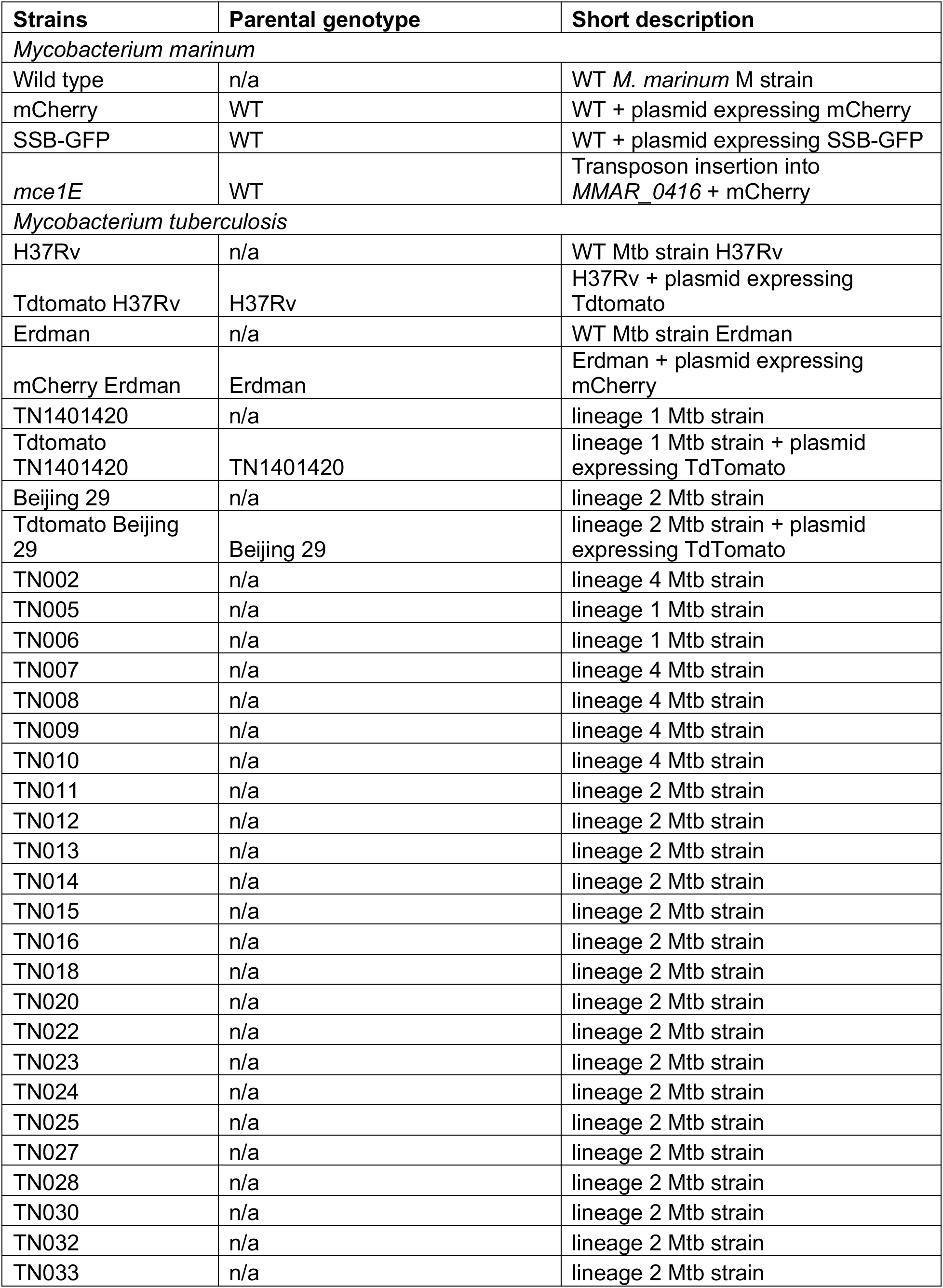
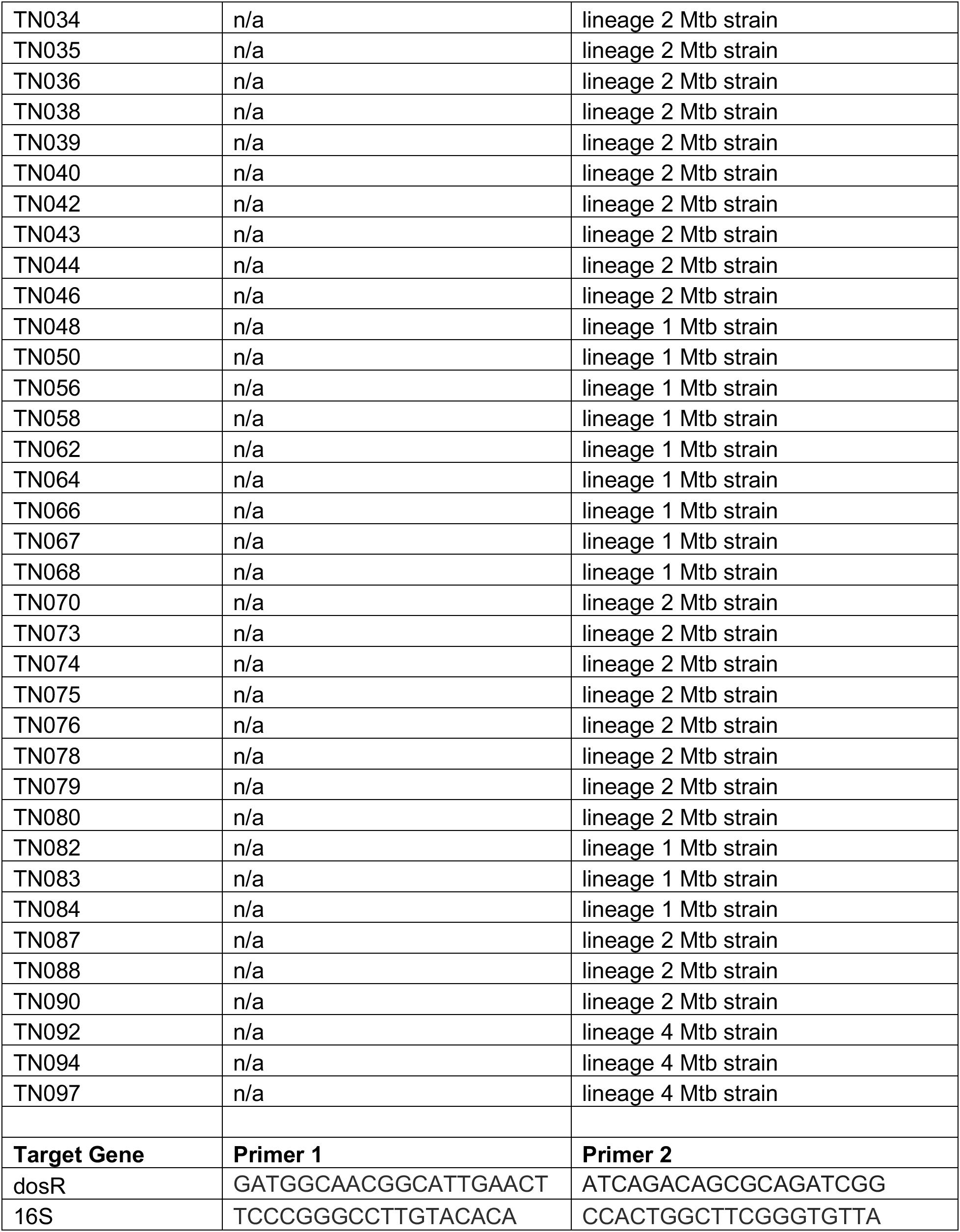
Strains and primers used in this work.

**Supplemental Video 1. Live imaging timelapse of *M. marinum* cultured in CellASIC flow chamber.** mCherry fluorescent WT *M. marinum* was cultured in a CellASIC flow chamber with 7H9^ADS^ media flowing through the system. Phase contrast images were taken every 15 minutes over the course of 17 hours.

**Supplemental Video 2. 3D reconstruction modeling of *M. marinum* cultured for 6 hours.** mCherry fluorescent WT *M. marinum* was cultured in 7H9^ADS^ for 6 hours and stained with BODIPY 493/503. 3D reconstruction modeling of SR-SIM images was performed using Imaris. Green structures represent BODIPY 493/503 signal and magenta structures represent mCherry signal.

**Supplemental Video 3. 3D reconstruction modeling of *M. marinum* cultured for 48 hours.** mCherry fluorescent WT *M. marinum* was cultured in 7H9^ADS^ for 48 hours and stained with BODIPY 493/503. 3D reconstruction modeling of SR-SIM images was performed using Imaris. Green structures represent BODIPY 493/503 signal and magenta structures represent mCherry signal.

**Supplemental Video 4. 3D reconstruction modeling of H37Rv cultured for 24 hours.** tdTomato fluorescent H37Rv was cultured in 7H9^OADC^ for 24 hours and stained with BODIPY 493/503. 3D reconstruction modeling of SR-SIM images was performed using Imaris. Green structures represent BODIPY 493/503 signal and magenta structures represent mCherry signal.

**Supplemental Video 5. 3D reconstruction modeling of Erdman cultured for 24 hours.** mCherry fluorescent Erdman was cultured in 7H9^OADC^ for 24 hours and stained with BODIPY 493/503. 3D reconstruction modeling of SR-SIM images was performed using Imaris. Green structures represent BODIPY 493/503 signal and magenta structures represent mCherry signal.

**Supplemental Video 6. 3D reconstruction modeling of L2 isolate cultured for 24 hours.** tdTomato fluorescent L2 isolate was cultured in 7H9^OADC^ for 24 hours and stained with BODIPY 493/503. 3D reconstruction modeling of SR-SIM images was performed using Imaris. Green structures represent BODIPY 493/503 signal and magenta structures represent mCherry signal.

**Supplemental Video 7. 3D reconstruction modeling of L1 isolate cultured for 24 hours.** tdTomato fluorescent L1 isolate was cultured in 7H9^OADC^ for 24 hours and stained with BODIPY 493/503. 3D reconstruction modeling of SR-SIM images was performed using Imaris. Green structures represent BODIPY 493/503 signal and magenta structures represent mCherry signal.

**Supplemental Data 1. RNA Seq DEG data of L2 compared to H37Rv, Erdman, L1.** H37Rv, Erdman, L1, and L2 strains were cultured in 7H9^OADC^ for 24 hours and sampled for RNA isolation and RNA Seq analysis. File represents all identified L2 DEG compared to H37Rv, Erdman, and L1 strains.

## Acknowledgements

The authors thank David Russell for the gift of the SSB-GFP plasmid; Midori Kato-Maeda for the gift of the L1 strain; William Jacobs for the gift of the L2 strain Beijing 29; Mary West at the UC Berkeley QB3 facility for assistance with confocal imaging; Azenta Life Sciences for RNA Sequencing and Analysis. We thank the members of the Stanley, Cox and Vance labs for helpful discussions, and Heran Darwin and Daisy Ji for reading a draft version of this manuscript. This work was supported by NIH grant AI113270 to SAS. DMF was supported in part by a Public Health Service Institutional Research Training Award NIH T32 GM007232.

## Author Contributions

Conceptualization, Methodology, and Analysis: DMF and SAS

Investigation: DMF, DS, AAS, and MK

Resources: NTT and JC

Writing - original draft: DMF

Writing – review & editing: DMF, SAS, JC

Funding Acquisition and Supervision: SAS

## References

[1] “WHO | Global Tuberculosis Report.” 2022.

[2] A. Cohen, V. D. Mathiasen, T. Schön, and C. Wejse, “The global prevalence of latent tuberculosis: a systematic review and meta-analysis,” Eur. Respir. J., vol. 54, no. 3, Sep. 2019.

[3] G. Gago, L. Diacovich, and H. Gramajo, “Lipid metabolism and its implication in mycobacteria-host interaction,” Curr. Opin. Microbiol., vol. 41, pp. 36–42, Feb. 2018.

[4] E. V. Nazarova et al., “Rv3723/LucA coordinates fatty acid and cholesterol uptake in Mycobacterium tuberculosis,” Elife, vol. 6, Jun. 2017.

[5] A. Gioffré et al., “Mutation in mce operons attenuates Mycobacterium tuberculosis virulence,” Microbes Infect., vol. 7, no. 3, pp. 325–334, Mar. 2005.

[6] D. G. Russell, P.-J. Cardona, M.-J. Kim, S. Allain, and F. Altare, “Foamy macrophages and the progression of the human tuberculosis granuloma,” Nat. Immunol., vol. 10, no. 9, pp. 943–948, Sep. 2009.

[7] I. Caire-Brändli et al., “Reversible lipid accumulation and associated division arrest of Mycobacterium avium in lipoprotein-induced foamy macrophages may resemble key events during latency and reactivation of tuberculosis,” Infect. Immun., vol. 82, no. 2, pp. 476–490, Feb. 2014.

[8] E. V. Nazarova, C. R. Montague, L. Huang, T. La, D. Russell, and B. C. VanderVen, “The genetic requirements of fatty acid import by Mycobacterium tuberculosis within macrophages,” Elife, vol. 8, Feb. 2019.

[9] K. Tauchi-Sato, S. Ozeki, T. Houjou, R. Taguchi, and T. Fujimoto, “The Surface of Lipid Droplets Is a Phospholipid Monolayer with a Unique Fatty Acid Composition*,” J. Biol. Chem., vol. 277, no. 46, pp. 44507–44512, Nov. 2002.

[10] J. Daniel et al., “Induction of a novel class of diacylglycerol acyltransferases and triacylglycerol accumulation in Mycobacterium tuberculosis as it goes into a dormancy-like state in culture,” J. Bacteriol., vol. 186, no. 15, pp. 5017–5030, Aug. 2004.

[11] A. P. Velázquez, T. Tatsuta, R. Ghillebert, I. Drescher, and M. Graef, “Lipid droplet-mediated ER homeostasis regulates autophagy and cell survival during starvation,” J. Cell Biol., vol. 212, no. 6, pp. 621–631, Mar. 2016.

[12] T. B. Nguyen et al., “DGAT1-Dependent Lipid Droplet Biogenesis Protects Mitochondrial Function during Starvation-Induced Autophagy,” Dev. Cell, vol. 42, no. 1, pp. 9–21.e5, Jul. 2017.

[13] R. J. Schulze, A. Sathyanarayan, and D. G. Mashek, “Breaking fat: The regulation and mechanisms of lipophagy,” Biochim. Biophys. Acta Mol. Cell Biol. Lipids, vol. 1862, no. 10 Pt B, pp. 1178–1187, Oct. 2017.

[14] E. R. Hinson and P. Cresswell, “The antiviral protein, viperin, localizes to lipid droplets via its N-terminal amphipathic α-helix,” Proceedings of the National Academy of Sciences, vol. 106, no. 48, pp. 20452–20457, 2009.

[15] P. T. Bozza, I. Bakker-Abreu, R. A. Navarro-Xavier, and C. Bandeira-Melo, “Lipid body function in eicosanoid synthesis: an update,” Prostaglandins Leukot. Essent. Fatty Acids, vol. 85, no. 5, pp. 205–213, Nov. 2011.

[16] J. Quillaguamán, H. Guzmán, D. Van-Thuoc, and R. Hatti-Kaul, “Synthesis and production of polyhydroxyalkanoates by halophiles: current potential and future prospects,” Appl. Microbiol. Biotechnol., vol. 85, no. 6, pp. 1687–1696, Feb. 2010.

[17] A. Steinbüchel and B. Füchtenbusch, “Bacterial and other biological systems for polyester production,” Trends Biotechnol., vol. 16, no. 10, pp. 419–427, Oct. 1998.

[18] H. M. Alvarez and A. Steinbüchel, “Triacylglycerols in prokaryotic microorganisms,” Appl. Microbiol. Biotechnol., vol. 60, no. 4, pp. 367–376, Dec. 2002.

[19] H. M. Alvarez, M. F. Souto, A. Viale, and O. H. Pucci, “Biosynthesis of fatty acids and triacylglycerols by 2,6,10,14-tetramethyl pentadecane-grown cells of Nocardia globerula 432,” FEMS Microbiol. Lett., vol. 200, no. 2, pp. 195–200, Jun. 2001.

[20] C. Zhang et al., “Bacterial lipid droplets bind to DNA via an intermediary protein that enhances survival under stress,” Nat. Commun., vol. 8, p. 15979, Jul. 2017.

[21] N. J. Garton et al., “Cytological and transcript analyses reveal fat and lazy persister-like bacilli in tuberculous sputum,” PLoS Med., vol. 5, no. 4, p. e75, Apr. 2008.

[22] H. L. Sheehan and F. Whitwell, “The staining of tubercle bacilli with Sudan black B,” J. Pathol. Bacteriol., vol. 61, no. 2, pp. 269–71, pl, Apr. 1949.

[23] L. G. Wayne and C. D. Sohaskey, “Nonreplicating persistence of mycobacterium tuberculosis,” Annu. Rev. Microbiol., vol. 55, pp. 139–163, 2001.

[24] L. E. Via et al., “Tuberculous granulomas are hypoxic in guinea pigs, rabbits, and nonhuman primates,” Infect. Immun., vol. 76, no. 6, pp. 2333–2340, Jun. 2008.

[25] K. L. Low et al., “Triacylglycerol utilization is required for regrowth of in vitro hypoxic nonreplicating Mycobacterium bovis bacillus Calmette-Guerin,” J. Bacteriol., vol. 191, no. 16, pp. 5037–5043, Aug. 2009.

[26] H. Ohno et al., “The effects of reactive nitrogen intermediates on gene expression in Mycobacterium tuberculosis,” Cell. Microbiol., vol. 5, no. 9, pp. 637–648, Sep. 2003.

[27] M. I. Voskuil et al., “Inhibition of respiration by nitric oxide induces a Mycobacterium tuberculosis dormancy program,” J. Exp. Med., vol. 198, no. 5, pp. 705–713, Sep. 2003.

[28] D. Raze et al., “Heparin-Binding Hemagglutinin Adhesin (HBHA) Is Involved in Intracytosolic Lipid Inclusions Formation in Mycobacteria,” Front. Microbiol., vol. 9, p. 2258, Sep. 2018.

[29] N. J. Garton, H. Christensen, D. E. Minnikin, R. A. Adegbola, and M. R. Barer, “Intracellular lipophilic inclusions of mycobacteria in vitro and in sputum,” Microbiology, vol. 148, no. Pt 10, pp. 2951–2958, Oct. 2002.

[30] S. Vijay et al., “Ultrastructural Analysis of Cell Envelope and Accumulation of Lipid Inclusions in Clinical Mycobacterium tuberculosis Isolates from Sputum, Oxidative Stress, and Iron Deficiency,” Front. Microbiol., vol. 8, p. 2681, 2017.

[31] P. Santucci et al., “Nitrogen deprivation induces triacylglycerol accumulation, drug tolerance and hypervirulence in mycobacteria,” Sci. Rep., vol. 9, no. 1, p. 8667, Jun. 2019.

[32] E. Iona, F. Giannoni, M. Pardini, L. Brunori, G. Orefici, and L. Fattorini, “Metronidazole plus rifampin sterilizes long-term dormant Mycobacterium tuberculosis,” Antimicrob. Agents Chemother., vol. 51, no. 4, pp. 1537–1540, Apr. 2007.

[33] M. O. Shleeva, Y. K. Kudykina, G. N. Vostroknutova, N. E. Suzina, A. L. Mulyukin, and A. S. Kaprelyants, “Dormant ovoid cells of Mycobacterium tuberculosis are formed in response to gradual external acidification,” Tuberculosis, vol. 91, no. 2, pp. 146–154, Mar. 2011.

[34] R. J. H. Hammond, V. O. Baron, K. Oravcova, S. Lipworth, and S. H. Gillespie, “Phenotypic resistance in mycobacteria: is it because I am old or fat that I resist you?,” J. Antimicrob. Chemother., vol. 70, no. 10, pp. 2823–2827, Jul. 2015.

[35] I. Mallick et al., “Intrabacterial lipid inclusions in mycobacteria: unexpected key players in survival and pathogenesis?,” FEMS Microbiol. Rev., vol. 45, no. 6, Nov. 2021.

[36] H.-D. Park et al., “Rv3133c/dosR is a transcription factor that mediates the hypoxic response of Mycobacterium tuberculosis,” Mol. Microbiol., vol. 48, no. 3, pp. 833–843, May 2003.

[37] T. D. Sirakova et al., “Identification of a diacylglycerol acyltransferase gene involved in accumulation of triacylglycerol in Mycobacterium tuberculosis under stress,” Microbiology, vol. 152, no. Pt 9, pp. 2717–2725, Sep. 2006.

[38] J. Daniel, H. Maamar, C. Deb, T. D. Sirakova, and P. E. Kolattukudy, “Mycobacterium tuberculosis uses host triacylglycerol to accumulate lipid droplets and acquires a dormancy- like phenotype in lipid-loaded macrophages,” PLoS Pathog., vol. 7, no. 6, p. e1002093, Jun. 2011.

[39] M. Knight, J. Braverman, K. Asfaha, K. Gronert, and S. Stanley, “Lipid droplet formation in Mycobacterium tuberculosis infected macrophages requires IFN-γ/HIF-1α signaling and supports host defense,” PLoS Pathog., vol. 14, no. 1, p. e1006874, Jan. 2018.

[40] J. Blanchette, M. Jaramillo, and M. Olivier, “Signalling events involved in interferon-gamma-inducible macrophage nitric oxide generation,” Immunology, vol. 108, no. 4, pp. 513–522, Apr. 2003.

[41] R. Bansal-Mutalik and H. Nikaido, “Mycobacterial outer membrane is a lipid bilayer and the inner membrane is unusually rich in diacyl phosphatidylinositol dimannosides,” Proc. Natl. Acad. Sci. U. S. A., vol. 111, no. 13, pp. 4958–4963, Apr. 2014.

[42] A. J. Martinot, et al., “Mycobacterial Metabolic Syndrome: LprG and Rv1410 Regulate Triacylglyceride Levels, Growth Rate and Virulence in Mycobacterium tuberculosis,” PLoS Pathog., vol. 12, no. 1, p. e1005351, Jan. 2016.

[43] S. Kurokawa et al., “Comparative genome analysis of fish and human isolates of Mycobacterium marinum,” Mar. Biotechnol., vol. 15, no. 5, pp. 596–605, Oct. 2013.

[44] T. Rogall, J. Wolters, T. Flohr, and E. C. Böttger, “Towards a phylogeny and definition of species at the molecular level within the genus Mycobacterium,” Int. J. Syst. Bacteriol., vol. 40, no. 4, pp. 323–330, Oct. 1990.

[45] T. Tønjum, D. B. Welty, E. Jantzen, and P. L. Small, “Differentiation of Mycobacterium ulcerans, M. marinum, and M. haemophilum: mapping of their relationships to M. tuberculosis by fatty acid profile analysis, DNA-DNA hybridization, and 16S rRNA gene sequence analysis,” J. Clin. Microbiol., vol. 36, no. 4, pp. 918–925, Apr. 1998.

[46] C. Barisch, P. Paschke, M. Hagedorn, M. Maniak, and T. Soldati, “Lipid droplet dynamics at early stages of Mycobacterium marinum infection in Dictyostelium,” Cell. Microbiol., vol. 17, no. 9, pp. 1332–1349, Sep. 2015.

[47] S. A. Stanley, S. Raghavan, W. W. Hwang, and J. S. Cox, “Acute infection and macrophage subversion by Mycobacterium tuberculosis require a specialized secretion system,” Proc. Natl. Acad. Sci. U. S. A., vol. 100, no. 22, pp. 13001–13006, Oct. 2003.

[48] Q. Zhang et al., “EsxA membrane-permeabilizing activity plays a key role in mycobacterial cytosolic translocation and virulence: effects of single-residue mutations at glutamine 5,” Sci. Rep., vol. 6, p. 32618, Sep. 2016.

[49] L. M. Stamm et al., “Mycobacterium marinum escapes from phagosomes and is propelled by actin-based motility,” J. Exp. Med., vol. 198, no. 9, pp. 1361–1368, Nov. 2003.

[50] M. Hagedorn, K. H. Rohde, D. G. Russell, and T. Soldati, “Infection by tubercular mycobacteria is spread by nonlytic ejection from their amoeba hosts,” Science, vol. 323, no. 5922, pp. 1729–1733, Mar. 2009.

[51] L. L. Listenberger and D. A. Brown, “Fluorescent detection of lipid droplets and associated proteins,” Curr. Protoc. Cell Biol., vol. Chapter 24, p. Unit 24.2, Jun. 2007.

[52] M. P. Weir, W. H. Langridge 3rd, and R. W. Walker, “Relationships between oleic acid uptake and lipid metabolism in Mycobacterium smegmatis,” Am. Rev. Respir. Dis., vol. 106, no. 3, pp. 450–457, Sep. 1972.

[53] C. McCarthy, “Utilization of palmitic acid by Mycobacterium avium,” Infect. Immun., vol. 4, no. 3, pp. 199–204, Sep. 1971.

[54] J. G. Rodríguez et al., “Global adaptation to a lipid environment triggers the dormancy-related phenotype of Mycobacterium tuberculosis,” MBio, vol. 5, no. 3, pp. e01125–14, May 2014.

[55] N. Sukumar, S. Tan, B. B. Aldridge, and D. G. Russell, “Exploitation of Mycobacterium tuberculosis reporter strains to probe the impact of vaccination at sites of infection,” PLoS Pathog., vol. 10, no. 9, p. e1004394, Sep. 2014.

[56] W. B. Schaefer and C. W. Lewis Jr, “Effect of oleic acid on growth and cell structure of mycobacteria,” J. Bacteriol., vol. 90, no. 5, pp. 1438–1447, Nov. 1965.

[57] A. S. Greenberg, J. J. Egan, S. A. Wek, N. B. Garty, E. J. Blanchette-Mackie, and C. Londos, “Perilipin, a major hormonally regulated adipocyte-specific phosphoprotein associated with the periphery of lipid storage droplets,” J. Biol. Chem., vol. 266, no. 17, pp. 11341–11346, Jun. 1991.

[58] M. Wältermann and A. Steinbüchel, “Neutral lipid bodies in prokaryotes: recent insights into structure, formation, and relationship to eukaryotic lipid depots,” J. Bacteriol., vol. 187, no. 11, pp. 3607–3619, Jun. 2005.

[59] M. Wältermann et al., “Mechanism of lipid-body formation in prokaryotes: how bacteria fatten up,” Mol. Microbiol., vol. 55, no. 3, pp. 750–763, Feb. 2005.

[60] M. Belton et al., “Hypoxia and tissue destruction in pulmonary TB,” Thorax, vol. 71, no. 12, pp. 1145–1153, Dec. 2016.

[61] L. E. Swaim, L. E. Connolly, H. E. Volkman, O. Humbert, D. E. Born, and L. Ramakrishnan, “Mycobacterium marinum infection of adult zebrafish causes caseating granulomatous tuberculosis and is moderated by adaptive immunity,” Infect. Immun., vol. 74, no. 11, pp. 6108–6117, Nov. 2006.

[62] S. Huang, W. Zhou, W. Tang, Y. Zhang, Y. Hu, and S. Chen, “Genome-scale analyses of transcriptional start sites in Mycobacterium marinum under normoxic and hypoxic conditions,” BMC Genomics, vol. 22, no. 1, p. 235, Apr. 2021.

[63] L. G. Wayne and L. G. Hayes, “An in vitro model for sequential study of shiftdown of Mycobacterium tuberculosis through two stages of nonreplicating persistence,” Infect. Immun., vol. 64, no. 6, pp. 2062–2069, Jun. 1996.

[64] K. H. Darwin, S. Ehrt, J.-C. Gutierrez-Ramos, N. Weich, and C. F. Nathan, “The proteasome of Mycobacterium tuberculosis is required for resistance to nitric oxide,” Science, vol. 302, no. 5652, pp. 1963–1966, Dec. 2003.

[65] R. Heintzmann and T. Huser, “Super-Resolution Structured Illumination Microscopy,” Chem. Rev., vol. 117, no. 23, pp. 13890–13908, Dec. 2017.

[66] N. Jacquier, V. Choudhary, M. Mari, A. Toulmay, F. Reggiori, and R. Schneiter, “Lipid droplets are functionally connected to the endoplasmic reticulum in Saccharomyces cerevisiae,” J. Cell Sci., vol. 124, no. Pt 14, pp. 2424–2437, Jul. 2011.

[67] F. Wilfling et al., “Triacylglycerol synthesis enzymes mediate lipid droplet growth by relocalizing from the ER to lipid droplets,” Dev. Cell, vol. 24, no. 4, pp. 384–399, Feb. 2013.

[68] M. B. Reed, S. Gagneux, K. Deriemer, P. M. Small, and C. E. Barry 3rd, “The W-Beijing lineage of Mycobacterium tuberculosis overproduces triglycerides and has the DosR dormancy regulon constitutively upregulated,” J. Bacteriol., vol. 189, no. 7, pp. 2583–2589, Apr. 2007.

[69] P. Domenech et al., “Unique Regulation of the DosR Regulon in the Beijing Lineage of Mycobacterium tuberculosis,” J. Bacteriol., vol. 199, no. 2, Jan. 2017.

[70] J. D. McKinney et al., “Persistence of Mycobacterium tuberculosis in macrophages and mice requires the glyoxylate shunt enzyme isocitrate lyase,” Nature, vol. 406, no. 6797, pp. 735–738, Aug. 2000.

[71] E. J. Muñoz-Elías and J. D. McKinney, “Mycobacterium tuberculosis isocitrate lyases 1 and 2 are jointly required for in vivo growth and virulence,” Nat. Med., vol. 11, no. 6, pp. 638– 644, Jun. 2005.

[72] E. J. Muñoz-Elías, A. M. Upton, J. Cherian, and J. D. McKinney, “Role of the methylcitrate cycle in Mycobacterium tuberculosis metabolism, intracellular growth, and virulence,” Mol. Microbiol., vol. 60, no. 5, pp. 1109–1122, Jun. 2006.

[73] J. A. Olzmann and P. Carvalho, “Dynamics and functions of lipid droplets,” Nat. Rev. Mol. Cell Biol., vol. 20, no. 3, pp. 137–155, Mar. 2019.

[74] K. M. Wilburn, R. A. Fieweger, and B. C. VanderVen, “Cholesterol and fatty acids grease the wheels of Mycobacterium tuberculosis pathogenesis,” Pathog. Dis., vol. 76, no. 2, Mar. 2018.

[75] O. Adeyo et al., “The yeast lipin orthologue Pah1p is important for biogenesis of lipid droplets,” J. Cell Biol., vol. 192, no. 6, pp. 1043–1055, Mar. 2011.

[76] W. Fei et al., “A Role for Phosphatidic Acid in the Formation of ‘Supersized’ Lipid Droplets,” PLoS Genet., vol. 7, no. 7, p. e1002201, Jul. 2011.

[77] J. R. Skinner et al., “Diacylglycerol enrichment of endoplasmic reticulum or lipid droplets recruits perilipin 3/TIP47 during lipid storage and mobilization,” J. Biol. Chem., vol. 284, no. 45, pp. 30941–30948, Nov. 2009.

[78] K. Ben M’barek, D. Ajjaji, A. Chorlay, S. Vanni, L. Forêt, and A. R. Thiam, “ER Membrane Phospholipids and Surface Tension Control Cellular Lipid Droplet Formation,” Dev. Cell, vol. 41, no. 6, pp. 591–604.e7, Jun. 2017.

[79] V. Choudhary et al., “Architecture of Lipid Droplets in Endoplasmic Reticulum Is Determined by Phospholipid Intrinsic Curvature,” Curr. Biol., vol. 28, no. 6, pp. 915–926.e9, Mar. 2018.

[80] J. Aronson, “Spontaneous Tuberculosis in Salt Water Fish,” J. Infect. Dis., vol. 39, no. 4, pp. 315–320, 1926.

[81] C. M. Sassetti and E. J. Rubin, “Genetic requirements for mycobacterial survival during infection,” Proc. Natl. Acad. Sci. U. S. A., vol. 100, no. 22, pp. 12989–12994, Oct. 2003.

[82] S.-H. Baek, A. H. Li, and C. M. Sassetti, “Metabolic regulation of mycobacterial growth and antibiotic sensitivity,” PLoS Biol., vol. 9, no. 5, p. e1001065, May 2011.

[83] L. Shi et al., “Carbon flux rerouting during Mycobacterium tuberculosis growth arrest,” Mol. Microbiol., vol. 99, no. 6, p. 1179, Mar. 2016.

[84] A. Blumenthal, C. Trujillo, S. Ehrt, and D. Schnappinger, “Simultaneous analysis of multiple Mycobacterium tuberculosis knockdown mutants in vitro and in vivo,” PLoS One, vol. 5, no. 12, p. e15667, Dec. 2010.

[85] T. Parish, “Electroporation of Mycobacteria,” Methods Mol. Biol., vol. 2314, pp. 273–284, 2021.

